# GPR34 regulates microglia state and loss-of-function rescues TREM2 metabolic dysfunction

**DOI:** 10.1101/2025.03.28.646038

**Authors:** Karthik Raju, David Joy, Sophia M. Guldberg, Kyla Germain, Neal S. Gould, Connie Ha, David Tatarakis, Jason Dugas, Sonnet S. Davis, Audrey G. Reeves, Shan V. Andrews, Tanya N. Weerakkody, Jung H. Suh, Gabrielly Lunkes de Melo, Ken Xiong, Yaima Lightfoot, Nicholas Propson, Thomas Sandmann, Gilbert Di Paolo, Joseph W. Lewcock, Jim Ray, Kathryn M. Monroe

**Author notes:** Lead contact/Corresponding author.

## Abstract

Microglia are implicated in modifying neurodegenerative disease risk in the central nervous system (CNS). GPR34 is a microglia-enriched G-protein coupled receptor that detects cytotoxic lipids upregulated in Alzheimer’s Disease (AD). Since dysregulated lipid metabolism occurs in disease, we hypothesized GPR34 could act with other lipid sensors, such as TREM2, to regulate microglial function. Here, we report that *GPR34* knockout (KO) rescues dysregulated cholesterol metabolism in *TREM2* KO iPSC-derived microglia (iMG) and alone promotes fatty acid catabolism without the proton leak observed in *TREM2* KO. Loss of *GPR34* downregulated ERK signaling, while its agonism promoted interaction with and activation of ERK. In healthy and amyloid mouse models, *Gpr34* KO accelerated the conversion of homeostatic microglia to disease-associated microglia (DAM) states. Additionally, in *Gpr34* KO amyloid mouse brain, the frequency of large plaques was increased compared to WT, indicating that *Gpr34* KO microglia may promote amyloid aggregation. Overall, our data suggest GPR34 as a therapeutic target for modulating microglial function to slow AD progression.

## INTRODUCTION

Microglia are a highly dynamic population of yolk-sac derived myeloid cells in the parenchyma of the CNS. They perform many innate immune functions vital to brain health, including surveillance, synaptic pruning, neurotrophic support, clearance of pathology, and a host of other activities to maintain physiological CNS function. During the neurodegenerative process in Alzheimer’s disease (AD), microglia transition from a homeostatic state to a spectrum of disease-responsive states associated with amyloid pathology and aging^1–5^. Since human genetics have implicated microglia as a key contributor to AD risk^6^, we sought to elucidate novel regulators of microglia state to identify potential new drug targets.

G protein-coupled receptors (GPCRs) are known to regulate macrophage transitions to more responsive states^7,8^. These receptors synchronize cellular responses to extracellular stimuli and play a vital role in the modulation of chemotaxis, phagocytosis, cell morphology, and other functions to coordinate an appropriately regulated immune response. For example, extracellular ADP and ATP agonize the purinergic GPCR P2RY12, reducing macrophage chemotaxis and polarization^9,10^. Additionally, CB_1_R/CB_2_R agonism reduces MCSF-dependent cell proliferation^11^. Given the sheer number of GPCRs identified in mouse and human genomes to date (>800), and their huge diversity in ligands, GPCRs represent a valuable druggable opportunity with demonstrated success in drugs targeting beta-adrenergic receptors (e.g., albuterol and salmeterol^12^) in the clinic for asthma, COPD, and other disorders.

GPR34 is a rhodopsin-like GPCR well-conserved (<90%) between mouse and human and is highly enriched in microglia^13,14^. Endogenous GPR34 ligands include lysophosphatidylserine (LysoPS), a lipid species upregulated in neuroinflammation^15^. GPR34 is expressed in microglia^16,17^, with genetic knockout (KO) associated with amoeboid morphology and altered phagocytosis in mice^18^. SiRNA-mediated knockdown of *GPR34* has been linked to reduced cytokine production^19^ in APP/PS1 mice. While these studies implicate GPR34 in disease-relevant pathways, little is known about the functional impact of GPR34 deficiency in microglia, or interactions with other master regulators of microglial cell states such as TREM2.

We therefore investigated the role of GPR34 in microglia via genetic deletion both in human iPSC-derived microglia (iMG) and in healthy aged and amyloid mouse models. In iMG, we examined the effects of *GPR34* KO alone and in the context of *TREM2* KO given the latter’s well-known associations with regulating microglia state and lipid biology^1,20–22^. *GPR34* KO corrected lipid metabolic defects in *TREM2* KO iMG by reducing cholesteryl ester (CE) accumulation upon myelin challenge. Moreover, *GPR34* KO alone was sufficient to reduce triglycerides (TGs) and enhance fatty acid oxidation (FAO) in iMG. We validated ERK activation as a key pathway downstream of GPR34 agonism. In both healthy and AD mouse models, *Gpr34* KO resulted in accelerated microglial transition from homeostatic to responsive DAM-like states, as evidenced by histological, morphological, transcriptional changes, and in reduced CEs, indicating *Gpr34*-dependent regulation of microglial state and lipid metabolism occurs in vivo. In the AD mouse model, these shifts in microglial responsiveness were even more pronounced in *Gpr34* KO, and were accompanied by an increase in the frequency of large plaques present in the mouse brain, suggesting a potential contribution to plaque compaction. Taken together, our data indicate GPR34 is a disease-relevant target that regulates microglial state, in which inhibition may be beneficial for slowing disease progression in AD.

## RESULTS

### *GPR34* KO reduces neurotoxic cholesteryl ester accumulation in microglia

Dysregulated cholesterol handling in microglia is *Trem2*-dependent^21,23,24^ and has been demonstrated in AD models^25,26^, as well as human disease^27^. Given the potential relevance for AD pathogenesis, we first examined the impact of *GPR34* KO on CE accumulation in CRISPR-edited iPSC-derived microglia (Figure S1A-S1C; Table S1) in the presence or absence of TREM2 after a 24-hour myelin challenge by performing LC/MS-based metabolomic and lipidomic analyses. *GPR34* KO was sufficient to reduce CE accumulation in most detected species by ≥50% in iMGs (Figure 1A). In addition, *GPR34* KO significantly (>50%) reduced CE species that accumulated in *TREM2* KO iMG, such as polyunsaturated fatty acid species CE(20:4), CE(20:5) and CE(22:6) (Figure 1A). Moreover, *GPR34* KO rescued several species of bis(monoacylglycerol)phosphate (BMP) dysregulated in *TREM2* KO (Figure 1A). These data show loss of GPR34 corrected *TREM2* KO-dependent lysosomal lipid dysregulation^28^. Both cholesterol sulfate and free cholesterol levels were unaffected (Figure 1A). To determine if this phenotype translated *in vivo*, we used fluorescence-activated cell sorting (FACS) to isolate microglia from *GPR34* KO and WT mouse brains and found that KO microglia exhibited reduced CEs in both young (4 month) and old (17 month) animals, with no impact to cholesterol sulfate and free cholesterol (Figure 1B). In parallel, we performed bulk RNA sequencing of WT and *GPR34* KO iMGs which revealed upregulation of genes associated with an oxysterol/LXR-responsive transcriptional program^29^ at baseline (buffer alone) (*FAS, ABCA1,* and *ABCG1*; Figure 1C; Table S2) or following a 24-hour myelin treatment (*SREBF1, LDLR,* and *FAS*; Figure 1C; Table S2). Other LXR-sensitive genes, including *APOE, LPL, APOC1, and MLXIPL*, were downregulated in the presence or absence of myelin (Figure 1C; Table S2). Taken together, these data implicate GPR34 signaling in cholesterol metabolism (Figure 1D).

**Figure 1.**
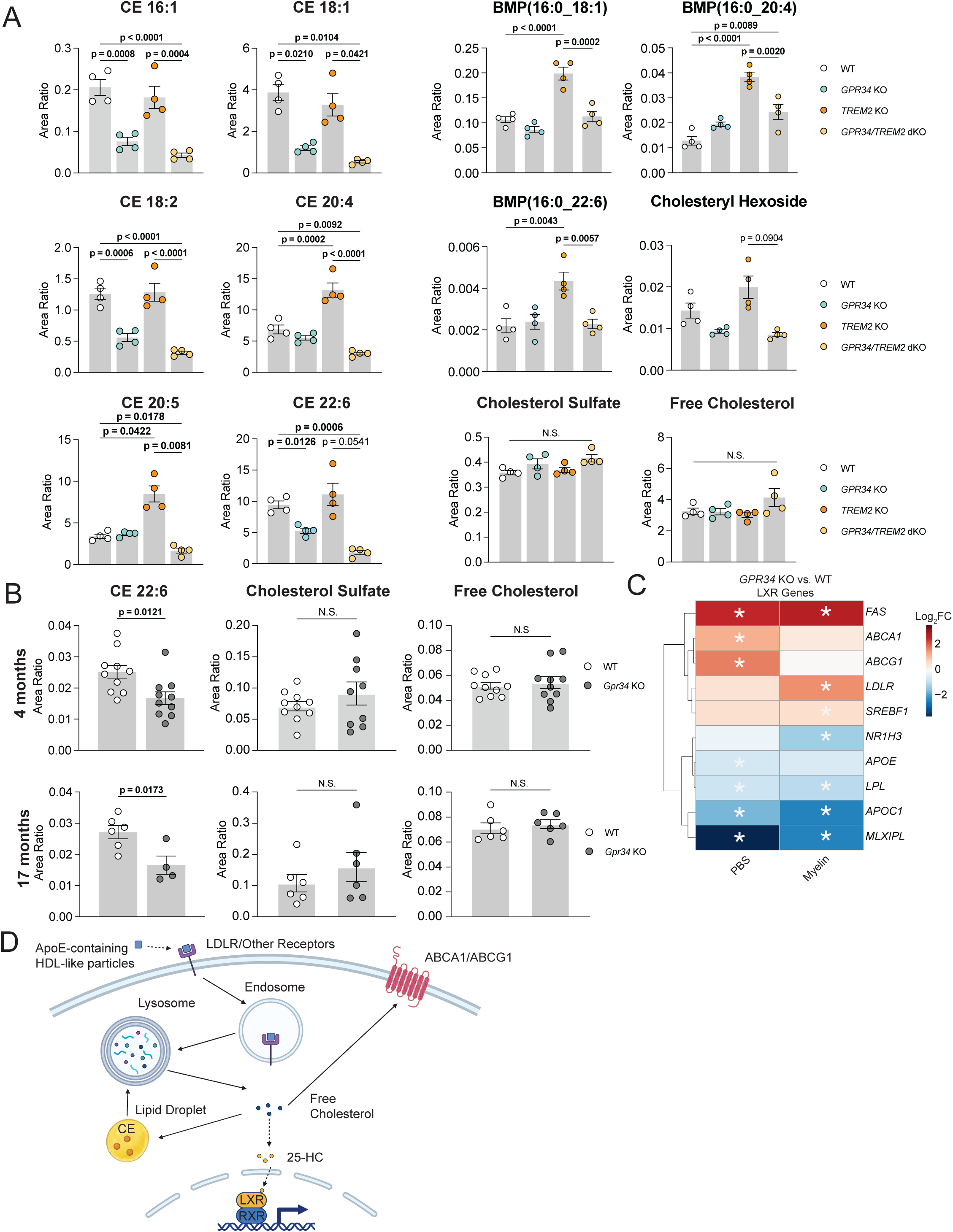
*GPR34* KO rescues neurotoxic cholesteryl ester accumulation alone and in *TREM2* KO iMG. **A)** LC-MS-based detection of relative abundance in microglial cholesteryl ester (CE), Bis(monoacylglycero)phosphate (BMP), cholesterol hexoside, cholesterol sulfate, and free cholesterol between WT, *GPR34* KO, *TREM2* KO, and *GPR34/TREM2* double KO (dKO) iMGs. Mean ± SEM (n=4 independent experiments), compared using one-way ANOVA followed by Sidak’s multiple comparisons test (CE) or Kruskal-Wallis multiple comparisons test (BMP). P-values < 0.05 are indicated in bold. **B)** LCMS-based lipidomic detection of relative abundance in CE, cholesterol sulfate, and free cholesterol within sorted microglia from WT and *Gpr34* KO mouse brain. Mean ± SEM (n = 4-10 animals/genotype), compared using an unpaired t-test. P-values < 0.05 are indicated in bold. **C)** Heatmap of differential gene expression (DGE) analysis of iMG bulk RNA-seq, shown as log_2_ fold change (Log_2_ FC), between *GPR34* KO and WT for LXR genes. Asterisks indicate adjusted p-value of < 0.1 across all hallmark gene sets. **D)** Pathway schematic illustrating cholesterol pathways in microglia.

### *GPR34* KO enhances mitochondrial oxidation of free fatty acids in microglia

Impairments in microglial bioenergetics have been described in mouse models of AD^30,31^, in TREM2-deficient mice, and in iMG models^21,30,32^. Given this precedent, we investigated the effects of *GPR34* KO alone or in combination with *TREM2* KO on FAO in iMGs. We observed elevated basal respiration in *GPR34* KO, *TREM2* KO, and *GPR34/TREM2* double KO (dKO) iMG by 50-80% relative to WT when using the fatty acid palmitate as the primary substrate (Figures 2A and 2B). However, in *TREM2* KO or dKO iMG, increased respiration was accompanied by ∼2-fold increase in proton leak (Figure 2C), while *GPR34* KO iMG showed no detectable impact on proton leak relative to WT iMG (Figure 2C). These data suggest that loss of *GPR34* enhances mitochondrial efficiency during fatty acid oxidation in contrast to *TREM2* KO, which exhibits decreased efficiency due to increased proton leak. Increased mitochondrial respiration in *GPR34* KO and *TREM2* KO iMG coincided with a 50-75% reduction in triglyceride levels relative to WT and 40-50% increase in beta-oxidation ratio^33^ after myelin challenge (Figures 2D and 2E), indicating that loss of either receptor drives catabolism of lipid energy sources. Given the functional increase in FAO present in *GPR34* KO relative to WT iMGs without elevated proton leak, it is not surprising that *GPR34* KO iMGs exhibited increased transcriptional expression of several genes associated with FAO, including rate-limiting enzymes *CPT1A, HSD17B4, ACADVL, EHHADH,* and *ACSL1* (Figure 2F)^34^. These data suggest that *GPR34* KO leads to increased metabolic efficiency in microglia for FAO without compromising mitochondrial function. In contrast, our data suggest that *TREM2* KO iMG likely upregulates respiration to compensate for significant proton leak from mitochondria.

**Figure 2.**
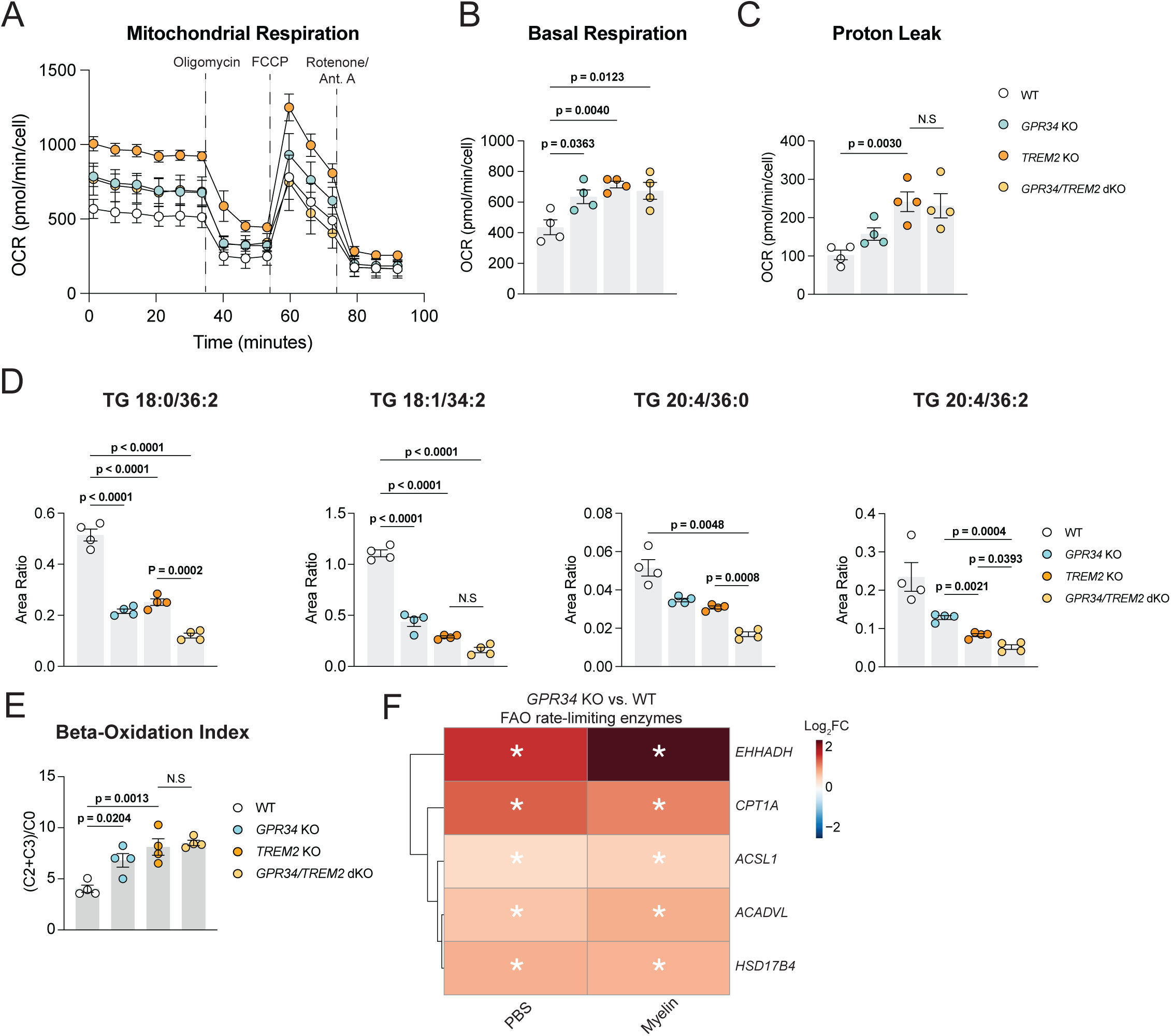
*GPR34* KO increased fatty acid catabolism without proton leak in iMGs. **A)** Representative plot of oxygen consumption in WT, *GPR34* KO, *TREM2* KO, and *GPR34/TREM2* double KO (dKO) iMGs upon palmitate treatment measured using the Seahorse Palmitate Oxidation assay. Mean ± SD (n=4 technical replicates). **B)** Basal respiration detected by Seahorse in all four iMG genotypes. Mean ± SEM (n=4 independent experiments), compared using one-way ANOVA followed by Tukey’s multiple comparisons test. P-values < 0.05 are indicated in bold. **C)** Proton leak detected in each genotype during the Seahorse assays described in A. Mean ± SEM (n=4 independent experiments), compared using one-way ANOVA followed by Sidak’s multiple comparisons test. P-values < 0.05 are indicated in bold. **D)** LCMS-based lipidomic detection of relative abundance in triacylglycerol/triglyceride species in all iMG genotypes 24h post myelin challenge. Mean ± SEM (n = 4 independent experiments), compared using one-way ANOVA followed by Sidak’s multiple comparisons test for 18:0/36:2 and 18:1/34:2, or Brown-Forsythe and Welch one-way ANOVA followed by Dunnett’s T3 multiple comparisons test for 20:4/36:0 and 20:4/36:2. P-values < 0.05 are indicated in bold. **E)** Beta-oxidation index after 24h myelin challenge using ratios of the sum of acetylcarnitine (C2) and propionylcarnitine (C3) to free carnitine (C0); metabolites detected by LC-MS. Mean ± SEM (n=4 independent experiments), compared using one-way ANOVA followed by Tukey’s multiple comparisons test. P-values < 0.05 are indicated in bold. **F)** Heatmap of differential gene expression (DGE) analysis of iMG bulk RNA-seq, shown as log_2_ fold change (Log_2_ FC), between *GPR34* KO and WT for rate-limiting FAO enzymes. Asterisks indicate adjusted p-value of < 0.1 across all hallmark gene sets.

### GPR34 agonism demonstrates temporal regulation of its interactome

To investigate the mechanisms by which GPR34 impacts cholesterol metabolism and mitochondrial function in microglia, we employed biotin ligase (BioID)-catalyzed proximal protein labeling^35,36^ to assess the kinetics of the GPR34 interactome following treatment with the M1 GPR34 agonist, a more stable analog of the native ligand lysophosphatidylserine^37^. Labeling was performed in phorbol-12-myristate-13-acetate (PMA)-differentiated U937 cells stably expressing GPR34 or a transmembrane control protein (TM Ctrl) fused to a C-terminal intracellular BioID to detect changes associated biotinylated proteins across several timepoints following treatment with agonist (Figure 3A). With this approach, we observed 80-200% enrichment of known G_α_ adaptor GNAS 0.5-10 minutes post M1-treatment, followed by detectable (>60%) enrichment of β-arrestin 2 (ARRB2) at 10-30 minutes in GPR34-BioID relative to TM Ctrl cells (p < 0.05 for each protein using a robust linear model in the Limma package in R; Figure 3B; Table S3). These data validated the utility of GPR34-BioID in mapping canonical GPCR signaling nodes via both initial G_α/β/γ_ and subsequent endosomal signaling, but surprisingly indicated GNAS as a primary G_α_ adaptor over GNAI, unlike previous descriptions in the literature for GPR34^37–40^.

**Figure 3.**
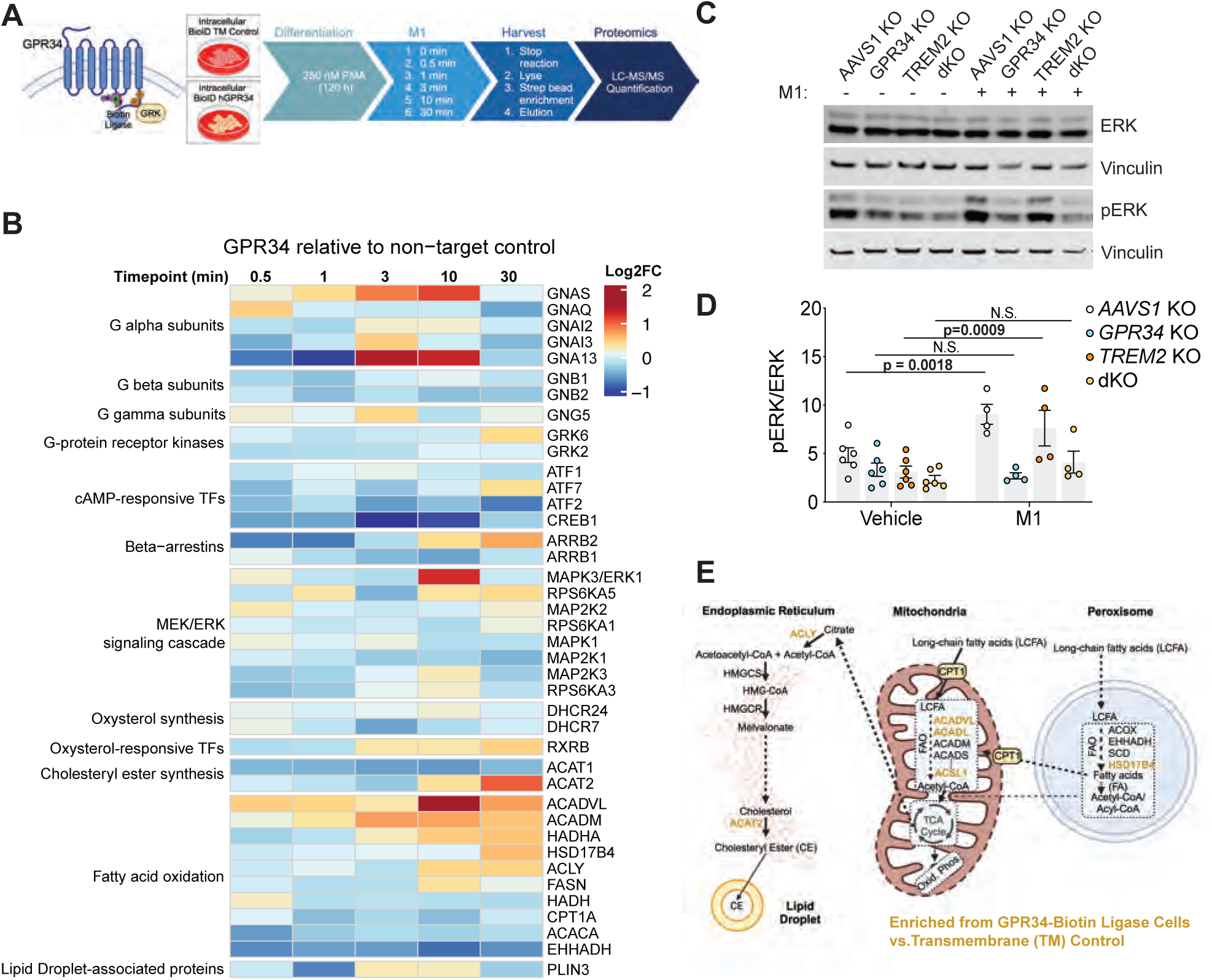
GPR34 activation demonstrates temporal regulation of its interactome. **A)** Schematic of GPR34 fused to biotin ligase (BioID) and experimental design for assessing the proximal interactome over time relative to a negative transmembrane control (TM Control) in stably expressing U937 cells. **B)** Heat map demonstrating mean ratios in the biotinylated protein fraction of GPR34-BioID vs. TM Control cells of known GPCR interactors and cholesterol metabolism, FAO, and lipid droplet proteins (n = 6 independent experiments). Protein levels at 0.5, 1, 3, 10, and 30 minutes post-M1 agonism were normalized to time 0, data expressed as Log_2_ fold-change ratio of GPR34-BioID vs. TM Ctrl. **C)** Representative Western blot of ERK and phospho-ERK (pERK) in iMGs CRISPR-edited to generate genetic deletion of *AAVS1* (non-targeting negative control), *GPR34*, *TREM2*, or double KO of both *GPR34* and *TREM2* (dKO). Vinculin serves as a loading control. **D)** Densitometry quantification of p-ERK/ERK ratio normalized to Vinculin in either vehicle- or M1-treated iMGs, with Mean ± SEM (n=4 independent experiments), compared using two-way ANOVA followed by Tukey’s multiple comparisons test. **E)** Pathway diagram showing enzymes enriched in the biotinylated fraction of GPR34-BioID cells relative to TM control (indicated in red) for cholesterol metabolism and endoplasmic reticulum- or mitochondrial-associated FAO.

Interestingly, β-arrestin levels after 10-30 minutes coincide with a transient ∼200% enrichment in MAPK3/ERK1 at 10 min (p < 0.05 using a robust linear model in the Limma package in R; Figure 3B; Table S3). To determine if this interaction was functionally relevant, we measured the phosphorylation status of ERK^41^ following M1 agonism in iMGs edited post differentiation via CRISPR with guide RNAs (gRNAs) to generate KOs against AAVS1 (as a CRISPR editing negative control), *GPR34*, *TREM2*, or both *GPR34* and *TREM2* as an orthogonal approach for investigating *GPR34*-dependent microglial biology (Figure S1D-S1G). M1 agonism resulted in >2-fold ERK activation in control *AAVS1* KO or *TREM2* KO iMGs which was ablated in *GPR34* KO or dKO iMG (Figures 3C-3D). These data suggest M1-induced interactions detected in GPR34-BioID U937 cells occur in an endogenous GPR34-expressing microglial model and contribute to upregulated ERK signaling. Lastly, we observed >20% enrichment in the CE synthesis enzyme ACAT2 and FAO enzymes HSD17B4, ACADVL, and ACADM at 10-30 min in GPR34-BioID cells (p < 0.05 for each protein using a robust linear model in the Limma package in R; Figure 3B; Table S3). These findings suggest that the interaction of GPR34 with ERK1/2 may be required for subsequent interactions with proteins involved in CE and FAO pathways and thus might be responsible for these phenotypes in *GPR34* KO iMG (Figures 1-2).

### *Gpr34* KO accelerates responsive microglial states accompanied by increased metabolic activity in vivo

To evaluate the impact of loss of *Gpr34* on microglial states in vivo, WT and *Gpr34* KO mice were crossed with an amyloid pathology model^42^ (*App*^SAA^) and assessed at 4, 8, and 17 months of age. CD11b^+^ CD45^+^ myeloid cells were isolated from forebrains of mice at the three timepoints for single cell transcriptional profiling (Figure 4, Figures S2-S4). Myeloid cells were annotated as coarse cell types (microglia, monocytes, or macrophages) before subclustering and annotating of microglia using a reference map based on 5 published datasets^22,43–47^ spanning both sexes and a variety of genotypes and ages (Method Details and Figures S2-S3). We detected 8 microglia subclusters and 2 peripheral myeloid populations expressing previously annotated marker genes (Figure 4A-4B). At 8 and 17 months, *Gpr34* KO microglia showed a significant reduction in homeostatic microglia and corresponding increase in responsive disease-associated microglia subclusters, including intermediate (DAM-1) and advanced (DAM-2) subpopulations (Figure 4C) in non-diseased mouse brains. Amyloid pathology accelerated the transition to DAM-1 and DAM-2 states compared to non-diseased brain, with *App*^SAA^ *GPR34* KO microglia increasingly allocated in the more terminal DAM-2 cluster at 8 and 17 months (Figure 4C) compared to their *App*^SAA^ counterparts.

**Figure 4.**
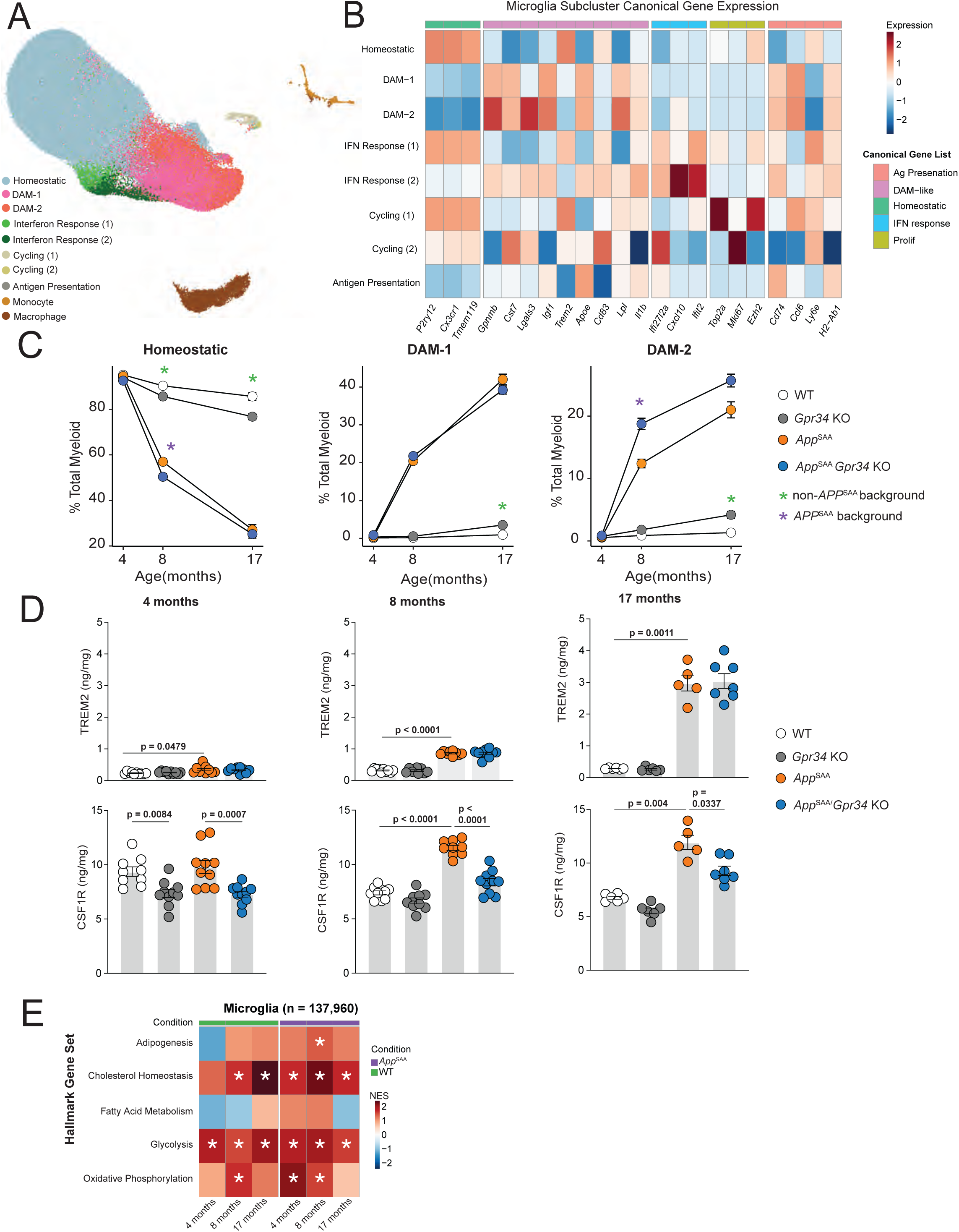
*Gpr34* KO accelerates responsive microglial states accompanied by increased metabolic activity in vivo. **A)** Integrated UMAP projection of scRNA-seq of ∼157,000 total myeloid cells from mice across entire study. Ten distinct subclusters of myeloid cells were identified; microglia subclusters were identified using cell label transfer via a reference map comprised of previously published mouse microglia scRNA-seq datasets (Method Details). **B)** Heatmap of canonical genes for each microglia subcluster. Expression represents average expression for each microglia subcluster scaled by column (gene). **C)** Microglial subcluster abundance as percentage of total myeloid cells for homeostatic (left), DAM-1 (middle), and DAM-2 (right) over time across all genotypes. Significance testing (t-test) was performed between *Gpr34* KO and WT in the *App*^SAA^ and non-*App*^SAA^ backgrounds, respectively, by timepoint (n=5-7 animals per genotype per age). Comparisons via t-test deemed significant after Bonferroni correction (p<(0.05/18) = 0.00278) are denoted via an asterisk (green = non-*App^SAA^* background, purple = *App^SAA^* background). Error bars represent standard error of the mean (SEM). **D)** TREM2 and CSF1R protein levels detected in whole brain across multiple ages. Comparisons involved one-way ANOVA with Brown-Forsythe and Welch tests, mean ± SEM (n=5-10 animals per genotype at each age, with p-values < 0.05 indicated in bold). **E)** Pseudo-bulked gene set enrichment analysis (GSEA) of all sorted microglia (n=137,960) from 4-, 8-, and 17-month-old WT and *Gpr3*4 KO mice on WT and *App*^SAA^ background, using the hallmark gene signatures collection. Heatmap is colored by normalized enrichment score (NES). Significant results are indicated with an asterisk and had an adjusted p-value < 0.1 across all hallmark gene sets.

To orthogonally assess markers of microglial state, we measured TREM2 and CSF1R protein levels in whole brain lysates. *Gpr34* KO had no effect on TREM2 levels in either WT or disease mice (Figure 4D), however, CSF1R levels were statistically significantly reduced with an effect size of 20-30% in *Gpr34* KO compared to WT in the pathology model across all ages (Figure 4D). Further gene set enrichment analyses (GSEA) of pseudo-bulked scRNA-seq data demonstrated *Gpr34* KO microglia upregulated genes annotated to cholesterol, glycolysis, and oxidative phosphorylation hallmark pathways regardless of age or disease (Figure 4E; Table S4), consistent with previous studies on DAM states in microglia^22^. In WT animals, transcriptional signatures associated with glycolysis and oxidative phosphorylation were transiently increased at 4-8 months in homeostatic microglia (Figure S4B; Table S4), with a subsequent elevation in DAM-2 microglia at 17 months in oxidative phosphorylation, adipogenesis, glycolysis, and cholesterol metabolism gene sets (Figure S4D; Table S4). In the presence of amyloid pathology, increased expression of glycolysis and oxidative phosphorylation gene signatures were consistent with observations in WT homeostatic microglia (Figures S4B; Table S4). However, in DAM-1 microglia at 8 and 17 months the signatures associated with glycolysis and cholesterol metabolism were elevated in *GPR34* KO relative to WT microglia (Figure S4C; Table S4). Taken together, single cell transcriptional profiling uncovered that *Gpr34* KO microglia increasingly transition to responsive microglia states relative to WT microglia in the presence or absence of amyloid pathology, with enriched metabolic gene signatures in cholesterol, glycolysis, and oxidative phosphorylation pathways.

### Loss of *Gpr34* drives microglial morphological changes and plaque compaction

To further investigate the impact of *Gpr34* in a disease context, we performed morphometric analyses of WT and *Gpr34* KO microglia in healthy and *App*^SAA^ mouse brain sections. *Gpr34* KO consistently resulted in an increased amoeboid microglial morphology compared to WT across multiple ages in *App*^SAA^ mice but not as robustly in non-disease conditions (Figures 5A-B and Figure S5A). *Gpr34* KO microglia in *App*^SAA^ mice displayed decreased ratio of cell surface area to volume, the percentage of total microglial cell volume at the cell surface (% surface voxels), and the condensation of microglial branching from ramified morphology to larger main branches (Figure 5A). Morphological alterations were accompanied by a 20-30% increase in total CD68 intensity after 4 months of age in the amyloid model (Figures 5A and 5B), which was not observed in healthy mice (Figure S5A).

**Figure 5.**
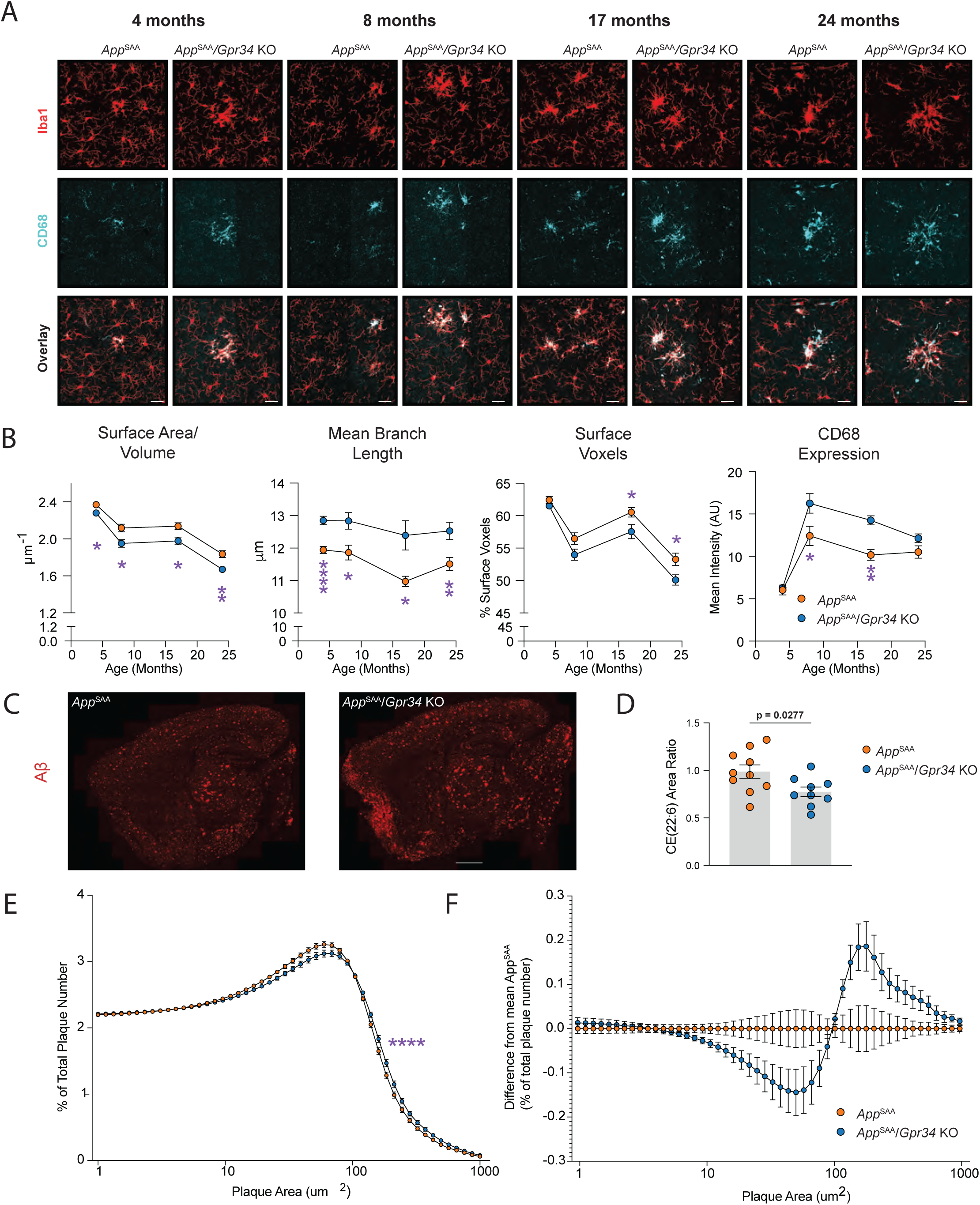
*Loss of Gpr34* drives microglial morphological changes and amyloid plaque compaction. **A)** Representative morphometric images of microglia detected by Iba1^+^ (red) and CD68 (cyan) in the cortex of *App*^SAA^; WT or *App*^SAA^; *Gpr34* KO mouse brain sections. Scale bars = 25 μm. **B)** Analysis of microglial morphology using surface area/volume, mean branch length, surface voxels, and CD68 levels. Mean ± SEM (n=5-10 animals per genotype at each age), with Student’s t-test used to compare between genotypes at each age (*p<0.05, **p<0.01, ****p<0.0001). **C)** Representative images of whole sagittal sections stained for beta-amyloid (red) in *App*^SAA^; WT or *App*^SAA^; *Gpr34* KO 24-month-old mouse brain sections. **D)** LCMS-based lipidomic detection of relative abundance in CE (22:6) content of sorted microglia from *App*^SAA^; WT or *App*^SAA^; *Gpr34* KO mouse brain. Mean ± SEM (n=4-10 animals/genotype), compared using an unpaired t-test. P-values < 0.05 indicated in bold. **E)** Frequency distribution of amyloid plaque sizes, mean ± SEM (n=8-10 animals per genotype), compared using the Kolmogorov-Smirnov test (****p< 0.0001). **F)** Frequency distribution of amyloid plaque sizes in WT vs *Gpr34* KO, normalized to *App*^SAA^ mouse brain sections.

Microglia are important contributors to amyloid plaque morphology and deposition in AD mouse models^48–50^. Based on our observations of *Gpr34* KO’s role in regulating microglia state and metabolism, we hypothesized *Gpr34* KO microglia may be better equipped to respond to amyloid pathology. Therefore, we assessed the impact of *Gpr34* KO on plaque size and distribution. We examined mice over a time course of 4, 8, 17, and 24 months (Figures 5C, 5E-5F, Figure S5B), and found only at 24 months *Gpr34* KO resulted in a subtle yet statistically significant (>10% effect size) decrease in the frequency of small plaques (<100 µm in diameter) in the cortex, with a concomitant increase in the number of plaques greater than this size (Figures 5E-5F). We did not observe any gross changes in amyloid load (Figure 5E). Moreover, we observed *Gpr34*-dependent decrease in microglial CEs in *App*^SAA^ mice (Figure 5D). Changes in CE were not observed in younger *App*^SAA^ mice (Figure S5C). Overall, this data reinforces that loss of *Gpr34* in microglia leads to increased microglial responsiveness and differential cholesterol metabolism in an amyloid pathology mouse model, correlating with a shift from smaller to larger amyloid plaques.

## DISCUSSION

This study examines the role of *GPR34* in regulating microglial biology to define the role of this GPCR in AD and determine whether modulation of GPR34 could be a viable therapeutic strategy. Here, we demonstrate GPR34 is an essential regulator of microglial homeostatic and responsive states in both WT and AD model mice.

Surprisingly, we found that loss of *GPR34* rescued lipid dysregulation associated with *TREM2* KO, by reducing previously described accumulation of CE and BMP species in microglia^21^. *GPR34* KO iMG also exhibited elevated basal rate of mitochondrial FAO, while lacking increased proton leak observed in *TREM2* KO iMG. Loss of *Gpr34* in vivo accelerated the transition of microglia from a homeostatic state to more responsive (DAM) states transcriptionally and histologically, with upregulation of gene signatures for cholesterol and fatty acid metabolism that persisted over time in the brains of healthy mice or in an amyloid model of AD (*App*^SAA^). *Gpr34* KO microglia demonstrated increased CD68 staining which coincided with a slight increase in the frequency of larger plaques present in 24-month-old mice. Together, these data indicate GPR34 is a key regulator of microglia whose endogenous function is to maintain a low metabolic and homeostatic state, likely to keep innate immune activity in check during healthy conditions. Given our disease-relevant findings in which GPR34 KO rescued *TREM2*-deficient metabolic dysfunction, GPR34 antagonism could be a potential therapeutic avenue to enhance microglial function in AD.

To elucidate GPR34 signaling and interactors, we conducted a detailed mechanistic analysis of small molecule-mediated GPR34 agonism in myeloid cells using proximity biotinylation. These studies demonstrated a temporal link between GPR34-dependent endosomal signaling, ERK activation, and subsequent interactions with cholesterol and mitochondrial cellular machinery. Moreover, our findings provide a potential molecular explanation for the phenotypes observed in *GPR34* KO iMG. Canonical GPCR studies and traditional GPCR therapeutic strategies have primarily focused on receptor signaling through G-protein subunits (G_αs_/G_αi_/G_αq_/G_β_/G_γ_)^51,52^. Based on heterogeneous expression studies in non-myeloid cells, GPR34 is thought to signal through G_αi_^37–40^. However, our proximity biotinylation studies of GPR34 signaling in myeloid U937 cells identified G_αs_/G_β_/G_γ_ as enriched G-protein effectors following M1 agonism^37^. Given that G_αi_ subunits showed no significant enrichment following GPR34 agonism in myeloid cells, this suggests that this receptor may utilize G_αs_ more than previously thought. We cannot completely rule out that our results may be explained by use of U937 as a myeloid cell model or a consequence of GPR34 overexpression. Therefore, validation in an endogenous cell system was important, and we confirmed ERK signaling as downstream pathway reduced in *GPR34* KO iMG consistent with previous findings^19^.

In recent years, GPCR-dependent endosomal signaling has emerged as another pathway through which these receptors modulate function. Through β-arrestin-mediated internalization via endosomes, GPCRs can potentiate alternative signaling cascades that have profound effects on protein function and cellular transcription^53,54^. β-arrestin depletion leads to reduced lipid accumulation in liver macrophages^55^, thereby providing a link between GPR34 and the mitochondrial phenotypes observed after its genetic deletion in myeloid cells. Many of these effects are dependent on endosomal GPCR signaling through ERK activation^56–58^, a pathway widely acknowledged to regulate cellular function^59^.

Alterations in ERK signaling have been associated with AD in several human and mouse studies. Increased phosphorylation of ERK and its substrates has been found in postmortem brain samples from AD patients^60^, with similar findings in multiple amyloid mouse models^19,61^. Several ERK antagonists, e.g. neflamapimod, are currently being evaluated in clinical trials for AD^62^. ERK activation in microglia has been proposed to act as a potentiator of amyloid pathology and neurodegeneration^63–65^. GPR34 may be a driver of dysfunction in microglia or inhibit protective responses in disease given that it is upregulated in AD human patients^66^ and *Gpr34* loss-of-function is associated with reduced ERK activation in APP/PS1 mice^19^, along with reduced amyloid accumulation in 5xFAD mice^66^. Our data provides the first evidence of microglia-specific ERK activation attributable to GPR34 agonism and offers a direct link between GPR34-dependent ERK activation and ERK-dependent effects on cholesterol metabolism^67^ and fatty acid catabolism^68,69^. Moreover, we describe mechanisms by which this potential novel target modulates microglial function in a more subtle manner compared to TREM2 antibody-mediated agonism^62,70^, which may avoid over-stimulated phenotypes associated with high TREM2-expressing microglia^6,71,72^. Since GPR34 is specifically expressed in microglia, targeting it could provide a cell-type specific way to suppress ERK signaling^62,70^. These findings indicate GPR34 inhibition is a novel therapeutic strategy to pursue either alone or in combination with amyloid-clearing approaches for AD.

## RESOURCE AVAILABILITY

### Lead Contact

Requests for further information and resources should be directed to and will be fulfilled by the lead contact, Kathryn Monroe.

### Materials Availability

All unique/stable reagents generated in this study are available from the lead contact with a completed materials transfer agreement.

### Data and Code Availability

Both bulk and single cell RNA-seq data generated during this study are available via the Gene Expression Omnibus (GEO; GSE293117). scRNAseq data for the datasets comprising the microglia state reference map are also available on GEO (GSE121654, GSE129788, GSE160523, GSE150358, GSE156183). Analysis objects, including that of the reference map, and code will be made available in Zenodo upon publication. For proteomic data from proximity biotinylation studies, all Bruker.d files and SpectroNaut processed results will be uploaded to a ProteomeXchange MASSIVE partner repository. Metabolomics data will be uploaded to the Metabolomics Workbench repository.

## Supporting information

GPR34-BioID Plasmid Sequence

TM Ctrl-BioID Plasmid Sequence

## ACKNOWLEDGEMENTS

We thank Lily Saraffha for help with flow cytometry. We thank Monroe lab members, the Discovery Biology group at Denali, contributors at MD Anderson in the the Neurodegeneration Consortium, as well as Tony Estrada and members of the Tenvie team for thoughtful discussions during the early stages of the project.

## AUTHOR CONTRIBUTIONS

K.R., K.M.M., Y.L.L., and J.R. conceived of the study idea and approaches. K.R., D.J., K.G., D.T., N.S.G, S.S.D., A.G.R., G.L.D.M., K.X., N.P., designed and performed experiments. K.R., D.J., S.M.G., K.G., D.T., C.H., S.V.A., J.D., T.S., analyzed and interpreted data. J.W.L, G.D.P, T.N.W., and S.V.A., edited the manuscript. K.R., K.G., K.M.M, S.M.G., and N.S.G. wrote the manuscript.

## SUPPLEMENTAL INFORMATION

**Document S1. Figures S1-S5, Table S1, and Prism Analyses’ File**

**Document S2. Sequencing Files for CRISPR-edited iMGs**

**Document S3: Table S3 and BioID construct sequences**

**Document S4. Tables S2 (full iMG bulk RNA-sequencing differential gene expression list, related to Figures 1C and 2F) and S4 (GSEA results for mouse scRNA-seq from bulk microglia and microglia subclusters, related to Figure 4)**

## SUPPLEMENTARY FIGURE LEGENDS

**Figure S1. CRISPR-mediated strategy for generation and validation of genetic KO of *GPR34* in human iPSC lines and acute KO in differentiated iMG.**

**A)** Flow cytometry scatterplots showing gating strategy for all cells, singlets, and iMG (CD11b^+^CD45^+^). **B)** Flow cytometry scatterplots showing iMG populations in WT, *GPR34* KO, *TREM2* KO, and dKO iMG lines, with percentage of singlet gate marked. **C)** Guide RNAs used to generate CRISPR-mediated *GPR34* KO*, TREM2* KO, and dKO lines. **D)** Guide RNAs used for acute CRISPR-mediated KO of *AAVS1*, *GPR34*, or *TREM2* in differentiated iMG**. E)** Western blot of iMG lysates 7 days after RNP nucleofection for acute KO, probed for TREM2 and Vinculin loading control. **F)** Distribution of deletion length within *GPR34* amplicon beginning 150 bp into locus, from gDNA collected 7 days after RNP nucleofection for acute KO. **G)** Integrative Genomics Viewer visualization of sequencing read alignments, and coverage plots of gDNA from F aligned to *GPR34* amplicon sequence. Each sample includes two tracks: the coverage plot (gray histogram) indicating sequencing depth across locus, and the aligned read (BAM) track, where individual reads are represented by horizontal gray bars. Black gaps within the read alignments indicate indels.

**Figure S2. Microglia reference map generation. A)** Schematic showing which published datasets were curated and how they were processed and integrated into a reference map. Experimental groups, sex, genotype, and group size are indicated for all studies in the data curation portion of the schematic. Conditions: Hammond: P30 = juvenile age wild-type, P100 = adulthood age wild-type, P540 = old age wild-type; Ximerakis: Young = 2-3 months of age wild-type, Old = 21-23 months of age wild-type; Huang: ADAM = A*pp*^SAA^ x *Axl*^-/-^*Mertk*^-/-^, AD = *App*^SAA^; Wang: CV/KO - IgG = 5XFAD x hTREM2 common variant treated wtih IgG, R47H/KO - IgG = 5XFAD x hTREM2 R47H variant treated with IgG, CV/KO - AL002c = 5XFAD x hTREM2 common variant treated with agonist hTREM2 antibody, R47H/KO - AL002c = 5XFAD x hTREM2 R47H variant treated with agonist hTREM2 antibody; Ellwanger: CV – IgG = 5XFAD x hTREM2 common variant treated with IgG, CV - hT2AB = 5XFAD x hTREM2 common variant treated with agonist hTREM2 antibody, KO – IgG = 5XFAD x mTrem2 knock-out treated with IgG, KO – hT2AB = 5XFAD x mTrem2 knock-out treated with agonist hTREM2 antibody, R47H – IgG = 5XFAD x hTREM2 R47H variant treated with IgG, R47H – hT2AB = 5XFAD x hTREM2 R47H variant treated with agonist hTREM2 antibody. **B)** Integrated UMAP of 30,707 total microglia from 5 published studies^43–47^. 8 subclusters were identified using unsupervised clustering. **C)** Heatmap of canonical genes used for cluster annotation for each microglia subcluster. Expression represents average expression for each microglia subcluster scaled by column (gene). **D)** Stacked bar plots of microglia subcluster composition for all samples integrated into the reference map. Each study is an individual plot and indicated by a different color (Ellwanger – blue; Hammond – purple; Huang – orange; Wang – green; Ximerakis – gray). Colors on the left indicate different conditions present in each respective study. **E)**. Stacked bar plots of cell subcluster composition for all samples integrated into the reference map faceted by genotype. Colors match those in D.

**Figure S3. Microglia reference map testing. A)** UMAPs of testing dataset with original, published cluster annotations faceted by genotype and treatment group^22^. **B)** UMAPs of testing dataset using reference map to transfer cell annotations faceted by genotype and treatment group. **C)** Stacked bar plot showing the relationship between the original cluster annotation and reference map annotations. **D-E)** Heatmap of canonical genes for original cluster annotations (D) or for reference map annotations (E). Expression represents average expression for each microglia subcluster scaled by column (gene).

**Figure S4. scRNA-seq cluster composition and GSEA analyses. A)** Stacked bar plots of cell subcluster composition for all scRNA-seq samples faceted by genotype (columns) and age (rows). **(B-D)** Gene set enrichment analyses of microglial subclusters from mouse brain. Pseudo-bulked gene set enrichment analyses (GSEAs) of microglia subclusters: homeostatic (B), DAM-1 (C), and DAM-2 (D) from 4-, 8-, and 17-month-old WT vs. *Gpr3*4 KO mice in non-*App*^SAA^ and *App*^SAA^ background, using the hallmark gene signatures collection. Heatmap is colored by normalized enrichment score (NES). Significant results are indicated with an asterisk and had an adjusted p-value < 0.1. n represents the number of cells in each cluster.

**Figure S5. Age-dependent assessment of microglial morphology in non-diseased mice, and both plaque size and microglial cholesteryl ester content in *App*^SAA^ mice.**

**A)** Representative morphometric images of microglia detected by Iba1^+^ (red) and CD68 (cyan) in the cortex of WT or *Gpr34* KO mouse brain sections. Analysis of microglial morphology using surface area/volume, mean branch length, surface voxels, and CD68 levels. Mean ± SEM (n=5-10 animals per genotype at each age), with Student’s t-test used to compare between genotypes at each age (*p< 0.05, ** p<0.01, *** p<0.001). Scale bars = 25 μm. **B)** Frequency distribution of raw amyloid plaque sizes from 4-, 8-, and 17-month-old *App*^SAA^; WT or *App*^SAA^; *Gpr34* KO mouse brain, mean ± SEM (n=8-10 animals per genotype), compared using the Kolmogorov-Smirnov test (****p< 0.0001), and corresponding frequency distributions of amyloid plaque sizes in WT vs *Gpr34* KO, normalized to *App*^SAA^ mouse brain sections. **C)** LCMS-based lipidomic detection of relative abundance in CE (22:6) content of sorted microglia from 4-, 8-, and 17-month-old *App*^SAA^; WT or *App*^SAA^; *Gpr34* KO mouse brain. Mean ± SEM (n=4-10 animals/genotype), compared using an unpaired t-test.

## STAR METHODS

## EXPERIMENTAL MODEL AND STUDY PARTICIPANT DETAILS

### Cell Lines

#### iPSC culture

iPSC-derived microglia (iMG) were differentiated from a human, female iPSC line (Thermo Fisher Scientific, A18945, hereafter referred to as iPSC line 1) using a previously published and validated protocol^23^. iPSCs were maintained on matrigel-coated plates (Corning, CB356238) according to the manufacturer’s specifications in mTeSR-plus (STEMCELL Technologies, 100-0276) or TeSR-E8 (STEMCELL Technologies, 05990). Culture medium was replenished every day with fresh medium. Cells were passaged every 5–7 days using Gentle cell dissociation (STEMCELL Technologies, 100-0485) or ReLeSR (STEMCELL Technologies, 100-0484) according to the manufacturer’s specifications. For all in vitro experiments, iPSCs were cultured in 5% O_2_, 5% CO_2_ at 37°C.

#### Hematopoietic and iMG Differentiation

iPSC line 1 was maintained in mTESR-Plus (STEMCELL Technologies, 100-0276) until seeding for differentiation. At ∼80% confluence hematopoietic differentiation was started as previously published^73^. On Day 4 cells were sorted for marker CD235a and replated on matrigel coated plates into Hematopoietic Kit Medium B (STEMCELL Technologies, 05310) containing 10µM ROCK inhibitor (ATCC, Y27632) at 500,000 cells per well of a 6-well plate. From this point onward all steps from STEMCELL Technologies protocol were followed to generate highly pure HPCs. Collected HPCs were confirmed to be >90% positive for marker CD43, seeded directly onto astrocyte feeder layers to generate iMG as previously reported^22^, and thereafter maintained in IMDM (Thermo Fisher Scientific, 31980030) supplemented with 10% FBS, 1X Penicillin/Strepyomycin, 20ng/mL hIL3 (Peprotech), 20ng/mL hGMCSF (Peprotech), and 20ng/mL hM-CSF (Peprotech) (hereafter referred to as microglia maintenance media).

#### Generation of *GPR34* and *TREM2* KO iPSC lines

All iPSC KO lines were generated from iPSC Line 1 using a CRISPR approach. *TREM2* KO line was generated in a prior study^21^ using a sgRNA sequence designed with the Broad Institute’s design tool (Figure S1C, Table S1). Briefly, iPSCs were maintained in mTESR-Plus (STEMCELL Technologies, 100-0276), combined with RNP complexes of sgRNA (ordered from IDT) and Cas9 (NEB, M0646M) and subjected to nucleofection using Lonza’s (V4XP-3032) P3 Primary Cell 4D-Nucleofector X Kit. *GPR34* KO and *GPR34*/*TREM2* dKO lines were generated by a contract research organization (Synthego) by introducing sgRNAs targeting exon 3 of GPR34 (cut location chr1:41,696,013) into WT or *TREM2* KO iPSCs (Figure S1C, Table S1). Due to lack of antibodies effective at detecting endogenous GPR34 in iMG, confirmation of KO was performed in differentiated iMG by gDNA sequencing (Document S2).

#### Validation of iMG differentiation by flow cytometry

iMG were transferred to V-bottom 96-well plates (50,000 cells/well) and washed once with fluorescence-activated cell sorting (FACS) buffer (1X HBSS, 25mM HEPES, pH 7.4, 2X Glutamax, 1% BSA, 1mM EDTA). Cells were resuspended in human Fc block (1:50, BioLegend, 422302) in FACS buffer. Following 15 minutes of blocking on ice, an equal volume of FACS buffer with 2x concentration of primary antibodies was added to the cells (CD11b-BV421, 1:50, BioLegend, 101235; CD45-APC/Cy7, 1:500, BD Biosciences, 561863, CD43-PE, 1:500, BioLegend, 343204). Cells were incubated for 30 minutes on ice and washed 3 times with FACS buffer before resuspension in FACS buffer. The samples were analyzed via flow cytometry using a BD FACSCanto II analyzer, the BD FACSDiva™ software, and the FlowJo software. The SSC-A and FSC-A parameters were used to set the cell gate, followed by the FSC-H and FSC-A parameters for the singlet gate (Figure S1A). The iMG gate was determined based on co-expression of the microglial markers, CD11b and CD45 (Figure S1A). iMG differentiation was not impacted by GPR34 or TREM2 KO (Figure S1B).

#### Acute CRISPR KO in differentiated iMG

iMG were differentiated from human iPSC Line 1 and validated as above and maintained in microglia maintenance media for at least 100 days. gRNA sequences targeting AAVS1, GPR34, or TREM2 were identified using the IDT CRISPR-Cas9 guide RNA design checker, ordered from IDT, and re-suspended in TE buffer at 100uM (Figure S1A, Table S1). Guides were complexed with Alt-R Sp Cas9 (IDT, 1081059) and subsequent RNP was delivered into iMG by nucleofection using Lonza’s P3 Primary Cell 4D-Nucleofector X Kit (V4SP3960). iMG were maintained for 24-h in Deepwell Protein LoBind plates (Eppendorf, 951032107) and thereafter collected and centrifuged to remove dead cells. Cells were plated in relevant assay plates and maintained for 9 days with half medium changes every 2 days. TREM2 KO was confirmed by protein loss via western blot analysis (Figure S1E) and GPR34 KO was validated by sequencing of gDNA as described below (Figure S1F-G).

#### Validation of *GPR34* KO by long read sequencing

gDNA was isolated from cells using the Quick-DNA^TM^ Miniprep Kit (Zymo Research, D3024) according to the manufacturers’ protocol. Primers flanking *GPR34* were designed and ordered from IDT (Table S1) and used for *GPR34* PCR amplification with Platinum^TM^ SuperFi II PCR Master Mixes according to the manufacturers’ protocol. *GPR34* PCR products were purified with QIAquick PCR purification Kit (Qiagen, 28106) and sequenced with a Oxford Nanopore Technologies MinION instrument at Plasmidaurus. Reads were aligned to the *GPR34* amplicon sequence via Minimap2^74^ (hg38, chrX:41695530-41697293). The resulting .bam files were filtered for reads spanning the intended deletion site. All surviving reads were then assessed for the number and length of deletions. To confirm an enrichment of deletions in the *GPR34* KO and dKO samples, but not the *AAVS1* KO or *TREM2* KO samples, we visually examined the alignments via the Integrative Genomics Viewer (IGV) software^75^ (Figure S1G) and plotted the distribution of deletion length in each sample (Figure S1F).

### Animals

*Gpr34* KO mice were generated at MD Anderson (KOMP allele ID Gpr34^tm1(KOMP)Vlcg^), and crossed in-house to non-diseased mice (Jackson Laboratory catalog no. 000664) or *App*^SAA^ mice (Jackson Laboratory catalog no. 034711) to generate all 4 genotypes used for the study, all maintained on a C57BL/6J genetic background. Only male animals were used for the study, since *Gpr34* is located on the X-chromosome. Mouse husbandry and experimental procedures were approved by the Denali Institutional Animal Care and Use Committee. Housing conditions included standard pellet food and water provided ad libitum, a 12-hour light/dark cycle at a temperature of 22 °C with maximal five mice per cage and cage replacement once per week with regular health monitoring.

## METHOD DETAILS

### RNA sequencing

#### Bulk RNA-seq sample and library preparation of iMG

WT, *GPR34* KO, *TREM2* KO, and *GPR34/TREM2* KO induced microglia (iMG) cultured in microglia maintenance media were treated with either PBS or myelin (25 µg/mL) for 24 hours. For each sample, RNA was extracted from a pool of two replicate wells, each seeded with 25,000–30,000 cells. A total of 32 RNA samples were collected, representing four genotypes, two treatments, and four replicates per condition. Each replicate was obtained from an independent harvest with a different seeding date, but all samples originated from the same differentiation batch. RNA extraction was performed using the RNeasy Plus Micro Kit (Qiagen, 74034), and RNA quality was assessed using the TapeStation High Sensitivity RNA ScreenTape (Agilent, 5067-5579).

Library preparation was conducted in a 96-well PCR plate using the QuantSeq 3’ mRNA-seq Library Prep Kit FWD for Illumina (Lexogen, A01173), incorporating the UMI second-strand synthesis module to identify and remove PCR duplicates, following the manufacturer’s protocol. Briefly, total RNA was used as input for oligo-dT priming during reverse transcription, followed by RNA removal. Unique Molecular Identifiers (UMIs) were introduced during second-strand synthesis, and cDNA was purified using magnetic beads, followed by 15 cycles of PCR with dual indexes and PCR purification with 0.8× magnetic beads.

Library quantity and quality were assessed using TapeStation High Sensitivity D1000 ScreenTape (Agilent, 5067-5584). Libraries were pooled in equimolar ratios, and the sequencing pool underwent an additional 0.8× magnetic bead purification to remove residual adapter contamination. Sequencing was performed on an Illumina NextSeq 5050 System High-Output Kit (SE, 75 bp, dual indexes) with a 5% PhiX spike-in, outsourced to SeqMatic (Fremont, CA, USA).

#### Bulk RNA-seq data mapping and preprocessing

Raw data (FASTQ files) were processed using nf-core/rnaseq v3.11.2 of the nf-core collection of workflows^76^ with GRCh38 genome sequences and gene annotations from Gencode version 42. To deduplicate unique molecular identifiers, the following umitools_bc_pattern was used: ^(?P.{6})(?P.{4}).*

To account for the fact that the QuantSeq 3’ mRNA-Seq library protocol only captures a 3’ tag from each transcript, the following custom argument was included in the nf-core/rnaseq workflow: extra_star_align_args: --alignIntronMax 1000000 -- alignIntronMin 20 --alignMatesGapMax 1000000 --alignSJoverhangMin 8 -- outFilterMismatchNmax 999 --outFilterType BySJout --outFilterMismatchNoverLmax 0.1 --clip3pAdapterSeq AAAAAAAA.

Similarly, the following custom arguments were passed to the salmon quantitation step of the workflow: extra_salmon_quant_args: --noLengthCorrection The nf-core/rnaseq workflow produces a SummarizedExperiment R object that is serialized to disk. To generate the processed data file, this object was read into an R session (using R 4.4.1), library sizes were estimated with the edgeR::normLibSizes function (version 4.4.2) using the TMM method and counts per million were returned with the edgeR::cpm function.

#### Bulk RNA-seq data analysis

Downstream analysis of bulk RNA-seq data was carried out in R (version 4.4.1). Differential expression was performed using the *limma/voom* (version 3.62.2) pipelines^77^. Genes were filtered based on expression across groups using *filterByExpr()* using default parameters. Differential expression was performed using *voomLmFit()* with default parameters and a model of “∼ 0 + group + batch”. Comparisons between GPR34KO and WT in both PBS and Myelin conditions were evaluated. Pathway enrichment was calculated through Gene Set Enrichment Analysis (GSEA) performed using the *fgsea*^78^ (version 1.32.2) R package using the Hallmark pathways^79,80^ as available via the Molecular Signatures Database (MSigDb; *msigdbr* version 7.5.1; Table S2)^80^. Visualizations were generated in R using the *ggplot2* (version 3.5.1) package.

#### Single cell RNA-seq microglia reference map generation

To infer microglia states/subtypes, we generated a reference map comprised of 5 publicly available datasets^43–47^ which included a variety of ages, genotypes, and perturbations. We focused specifically on mouse models and treatments used in the study of Alzheimer’s disease and wild-type mice to best build a map representing the microglia in our experimental data. For each publicly available dataset used in the reference map, data was read into a Seurat Object in R using *Seurat* (version 5.2.1)^81^. Each dataset was split by their experimental groups prior to normalizing counts using the log10 normalization with the “*NormalizeData*” function. “*FindVariableFeatures*” determined 2000 variable features using the “vst” method. Each dataset was scaled using default methods in “*ScaleData*” and PCs were generated using “*RunPCA*”. Each dataset was then integrated in the PC space using CCA integration with each experimental group as a batch. All datasets had standard filtering for nFeature, nCount, and percent mitochondrial reads. If applicable, data was subset to microglia only (Ximerakis dataset, Hammond dataset). If the dataset had cluster annotations provided, these were used as cluster annotations (Ellwanger dataset, Huang dataset). If the dataset did not have cluster annotations provided, “*RunUMAP*”, “*FindNeighbors*”, and “*FindClusters*” were run. Doublets were removed using *scDblFinder* (version 1.20.0)^82^. If applicable, clusters were annotated using canonical marker genes. Each dataset was then subsampled to decrease enrichment of homeostatic microglia to improve label transfer for rare microglia states downstream.

All datasets were then merged and pre-processing was performed again. Each dataset was then integrated in the PC space using CCA integration with each dataset/study as a batch. “*RunUMAP*”, “*FindNeighbors*”, and “*FindClusters*” were run to generate integrated clusters. These clusters were annotated using common microglia marker genes (Figure S2). Visualizations were generated using *ggplot2* (version 3.5.1) and *pheatmap* (version 1.0.12) packages.

Map label validation was performed by testing cell label transfer onto a publicly available dataset^22^ and comparing the mapped cell labels to the original published cluster annotations (Figure S3A-B). The original cluster annotations were compared to the reference map generated annotations (Figure S3C). Original clusters 1, 3, 4, and 5 mapped to clusters very similar to their original identity. Cluster 2 was split primarily between DAM-2 and Cycling (1). Further investigation into expression of canonical genes indicates that Cluster 2 expresses some DAM-like genes and some proliferative genes (Figure S3D). Upon reference mapping, proliferative genes are extremely specific for Cycling (1) and Cycling (2) clusters indicating increased granularity achieved using the microglia reference map compared to unsupervised clustering/manual annotation (Figure S3E).

#### Single cell RNA-seq sample and library preparation

Three cohorts of mice, aged 4 months, 8 months, and 17 months were sacrificed for single-cell RNA-seq analysis, with 23-24 mice per age group, totaling 71 animals. Each cohort contained four genotypes: *Gpr34* WT/*App* WT, *Gpr34* WT/ *App*^SAA^, *Gpr34* KO/ *App* WT, and *Gpr34* KO/ *App*^SAA^. Sample preparation for the 4-month cohort was divided into three batches of eight animals each, with two replicates per genotype per batch. For the 8-month and 17-month cohorts, samples were processed in two batches of 12 animals each, with three replicates per genotype per batch.

Whole forebrains were dissected out for each animal (excluding the olfactory bulb and cerebellum) and dissociated into a single-cell suspension following the Miltenyi Adult Mouse Brain Dissociation protocol (Miltenyi, 130-107-677) with modifications. Anisomycin (2 µM final concentration), Actinomycin D (5 µM final concentration), and Hyaluronidase Type I-S (5,600 U/mL final concentration) were added to the Miltenyi Enzyme Mix 1 to facilitate extracellular matrix breakdown and inhibit transcriptional and translational activity. Single-cell suspensions were co-labeled with anti-mouse Fc block, CD45, CD11b, CD44, and Ly6G (each at a 1:100 dilution) for 30 minutes on ice (see Key Resources Table) in 0.5% BSA/25 mM HEPES pH 7.4/0.5 mM EDTA in 1x HBSS. Following a post-FACS labeling wash, 0.5–1 million cells per sample were labeled with individual Cell Multiplexing Oligos (CMO, 10X Genomics, 1000261) per the manufacturer’s protocol, substituting the PBS wash buffer with NbActiv-1 + 1% BSA. Each batch contained either 8 or 12 CMO-tagged samples, which were counted using a hemocytometer and pooled at an equal cell number ratio. Next, we isolated CD11b+CD45-Ly6G-microglia from the cell pool by FACS and loaded them onto a Chromium microfluidic chip G, following the Chromium NextGEMS Single Cell 3’ v3.1 Cell Multiplexing RevB User Guide (10X Genomics, CG000388), targeting an estimated capture of 3,000 microglia per animal.

After reverse transcription, GEM purification, and cDNA amplification, the amplified material was split into two fractions: 3’ Gene Expression (GEX) DNA and Cell Multiplexing (CMO) DNA, using a dual-side size selection bead purification. Five microliters of the CMO DNA was PCR-amplified to introduce Illumina sequencing indices, while one-quarter of the GEX DNA was used for library preparation following the user guide. For each cohort, GEX libraries were pooled in equimolar amounts and subjected to a QC sequencing run on an Illumina MiSeq to determine the number of captured cells. Based on the MiSeq results, the libraries were re-pooled along with the CMO libraries, and sequenced on either an Illumina NovaSeq X Series or NovaSeq 6000 instrument (28 × 10 × 10 × 90 cycles) at SeqMatic (Fremont, CA, USA), targeting a read depth of 50,000 reads per cell for mRNA transcripts and 5,000 reads per cell for demultiplexing CellPlex oligos.

#### Single cell RNA-seq alignment and data analysis

Raw reads were aligned to the mouse genome (mm39) using cellranger multi (10X Genomics, version 9.0). Downstream analyses were performed in R (version 4.4.1) using the *Seurat* package (version 5.2.1)^81^. Cells with < 1000 features or > 5000 features or > 3% mitochondrial reads were excluded from downstream analysis. Counts were log10 transformed and normalized with the “*NormalizeData*” function. “*FindVariableFeatures*” determined 2000 variable features using the “vst” method. Data was scaled using default parameters of the “*ScaleData*” function, and Principal Components (PCs) were obtained using “*RunPCA*”. Data was then integrated in the PC space using harmony integration with each timepoint as a batch^83^. A UMAP embedding was generated after integration from the first 30 most significant PCs using “*RunUMAP*”. Features from the first 30 most significant PCs were used for unsupervised louvain clustering in the “*FindNeighbors*” and “*FindClusters*” functions with a resolution of 0.5. Doublets were removed using *scDblFinder* (version1.20.0)^82^. Non-myeloid cells were removed from the data using common marker genes. After subsetting to myeloid cells, data was preprocessed again now using a resolution of 0.3 for clustering.

The previously described reference map was used to assign cell labels to the microglia populations we obtained in our study^84^. Anchors were found using “*FindTransferAnchors*” and predictions were mapped using “*TransferData*”. These annotations were used for all downstream analyses.

Data was pseudobulked using sample ID and cell type (either all microglia or individual microglia subclusters) and using “*aggregateAcrossCells*” from the scuttle package (version 1.16.0). Pseudobulk differential expression analyses comparing GPR34KO to WT in non-*App*^SAA^ and *App*^SAA^ backgrounds, respectively, were carried out using *limma/voom* (version 3.62.2)^77^. GSEA analyses were carried out with fgsea (version 1.32.2)^78^. GSEA was conducted across all microglia and within microglia subclusters; for the latter analyses were only conducted if there were >5,000 cells in that state per group (Table S4). Hallmark gene sets were obtained from the Molecular Signatures Database (MSigDB; *msigdbr* version 7.5.1)^79,80^. Visualizations were generated using *ggplot2* (version 3.5.1) and *pheatmap* (version 1.0.12) packages.

### Lipidomics

#### Sample preparation for LC-MS

iMGs were cultured in microglia maintenance media before plating at a density of 25K cells/well (3 wells per sample, each sample consisting of one treatment + genotype) on a PDL-coated 96 well plate in fresh microglia maintenance media. Following overnight incubation at 37C, cells were switched to microglia maintenance media in the presence of PBS vehicle or 25 µg/ml myelin^21^ for 24 hours. After treatment, cells were gently washed twice in ice-cold PBS and harvested in 50 ul/well of MS grade methanol + internal standards. For lipid and metabolite extraction, samples were gently vortexed, and volumes were adjusted to 100μl with MS grade H2O, vortexed again, and transferred to 1.5 ml Eppendorf Protein LoBind tubes on ice). 100μl tert-butyl methyl ether (MTBE) was added per sample, followed by a brief vortex period and then centrifugation at 21,000 g for 10 min at 4C. The two phases generated by centrifugation were separated. Each phase (top: non-polar lipids, bottom: polar metabolites) was transferred to glass vials and dried overnight using a Genevac EZ3. Non-polar lipids were resuspended in MS grade methanol and polar metabolites were resuspended in a 9:1 methanol:water mixture. Four biological replicates were collected per treatment + genotype. For sorted microglia from mouse brain, 20K-100K microglia per brain were sorted directly into 400 ul of MS-grade methanol + internal standards. Samples were adjusted up to 600 ul with H2O, with 600 ul of tert-butyl methyl ether then added to each sample. Samples were then processed the same way as iMGs, with each sample representing an independent biological replicate. Relative quantification of lipids and metabolites were performed as previously described^85^.

### Proteomics

#### Generation of Plasmid Constructs

Plasmid cDNA constructs encoding human *GPR34* (Horizon Discovery catalog no. MHS6278-202841295) and human *EGFR* (Origene catalog # SC101205) were purchased and used as templates for generating BioID fusion constructs. All constructs were generated using NEB Hifi Assembly (New England Biolabs, catalog no. E2621) and SuperFi II PCR Master Mix (Thermo Fisher, catalog no. 12368010), according to manufacturer’s protocol. For GPR34-BioID, the coding sequence corresponding to a.a. 1-381 (Uniprot accession # Q9UPC5) in human *GPR34* was amplified by PCR from the previously indicated Horizon Discovery plasmid template and purified. Independently, a 2xGGGGS linker followed by a modified ultraID sequence^36^ was synthesized as a gBlock (IDT) and inserted in-frame after *GPR34* a.a. 1-381 in the C-terminus of the GPCR, all upstream of the IRES sequence in the pLVX-IRES-Puro Shuttle vector (Takara, catalog no. 632183). Downstream of the IRES, an mClover3^86^ -P2A sequence was synthesized as a gBlock (IDT) and fused in-frame to the N-terminus of the puromycin resistance gene. For the TM Control plasmid, the same sequence downstream of the IRES was used as the GPR34-BioID construct. For the gene upstream of IRES, the human *EGFR* signal peptide (a.a. 1-24, Uniprot accession # P00553) was amplified by PCR from the indicated OriGene template and assembled in-frame with a consecutive Myc/HA/FLAG epitope tag, followed by amino acids. 25-668 of human *EGFR* and a 2xGGGGS linker with a modified ultraID. Plasmid maps are included in Document S4.

#### Generation of U937 stable cell lines and sample preparation for proximity biotinylation studies

U937 cells were purchased from ATCC (catalog no. CRL-1593.2), and maintained in 10% fetal bovine serum (Avantor, catalog no. 97068-91) in RPMI-1640 base medium (Thermo Fisher, catalog no. 22400089) when not being used for proximity biotinylation experiments. Lentivirus expressing either GPR34-BioID or TM Ctrl-BioID were generated according to manufacturer’s protocol (Mirus Bio, catalog no. MIR6605) from Lenti-X HEK-293T cells (Takara, catalog no. 632180), and enriched from supernatant using Lenti-X Concentrator (Takara, catalog no. 631232) according to manufacturer’s protocol. Lentiviruses were then used to transduce U937 cells in the presence of LentiBlast Premium according to manufacturer’s protocol (Oz Biosciences, catalog no. LBPX500) for 3 days, and stable cells were expanded using puromycin selection at a concentration of 2 ug/ml for 14 days. For proximity biotinylation experiments, stable cells were maintained for 5 days hours in 10% dialyzed FBS (Thermo Fisher catalog no. A3382001)/0.25 µM 12-O-tetradecanoylphorbol-13-acetate (TPA; Cell Signaling Technology catalog no. 4174S) in RPMI-1640 base media to both differentiate cells to a more macrophage-like state^87^ and to ensure uniform media biotin concentration. Cells were then washed twice and equilibrated in base RPMI-1640 media on the day of experiment for at least 2 hours at 37°C. GPR34 or TM control cells were then treated with or without GPR34 agonist M1^37^ (custom synthesis by Wuxi) at a concentration of 25 nM for 0.5, 1, 3, 10, or 30 minutes at 37°C, washed twice in ice-cold PBS, and harvested in RIPA buffer (Teknova, catalog no. R3792). ^37^Lysates were incubated with NanoLINK streptavidin-coated magnetic beads (Vector Biolabs, catalog no. M1002) overnight at 4°C. Beads were washed twice with 1% SDS (Thermo Fisher, catalog no. 24730020) in Tris-buffered saline (TBS), twice with RIPA buffer, and twice with TBS (Teknova, catalog no. T9545), prior to storage as a dry bead pellet prior to sample processing for proteomic analysis.

#### Sample preparation for proteomics

Protein was eluted from streptavidin beads using 2x 100 uL hexafluoroisopropanol (HFIP). The pooled HFIP eluates were dried in a speedvac and then resuspended in 50 uL PBS + 5% SDS for processing via S-trap (Protifi) 96-well plate format according to manufacturer’s protocol. Briefly, samples were reduced with 15 mM TCEP at 60°C for 30 mins and 1000 rpm. 15mM Iodoacetamide was added and incubated in the dark for 1 hr. Samples were then acidified to 1.2% phosphoric acid and mixed with 7 volumes of S-trap binding buffer (50 mM triethylammonium bicarbonate (TEAB), 90% methanol pH 7.2). Samples were loaded on the S-trap plate at 1500xg for 2 min, then washed 3x with 200 uL S-trap binding buffer. Trypsin was added at 1:30 in 100 mM TEAB and incubated at 37°C overnight. Samples were eluted in 80 uL of TEAB, 0.2% formic acid, then 0.2% formic acid/50% acetonitrile and dried in the speedvac. Samples were resuspended in 0.1% formic acid, peptide was measured by nanodrop and adjusted to 0.5 ug/uL. Analysis used an Agilent 1290 fitted with split-flow nanoflow adapter coupled to a Bruker timsTOF Pro 2. Peptide (0.5 ug) was loaded on an Aurora Ultimate column 25cm x 75um ID with 1.7um (IonOpticks) with the flowrate set to approximately 300 nl/min at 98% mobile phase A (0.1% formic acid in water) and 2% mobile phase B (0.1% formic acid in acetonitrile) with a gradient of 22% B at 90 mins, 37% B at 105 min, 80% B from 115-120 min then reconditioning at 2% B between 120-145 min. The MS was set to 100 ms ramp time with a duty cycle of 100%, dia-PASEF settings were set to a mass range of 300-1400 Da with an Ion mobility range of 0.70-1.40 1/K_0_ and collision energy scaling from 20eV at 0.60 1/K_0_ to 65 eV at 1.60 1/K_0_. DIA windowing was determined based on prior runs using py_diAID^88^. Raw data was analyzed using Spectronaut version 19 default settings (Biognosys). Additional analysis was done using custom scripts in R.

#### Western blot analysis of ERK signaling

Mature iMGs were subjected to acute CRISPR KO as described above, seeded (40,000 cells/well) on PDL-coated 96-well microplates (Ibidi, 89606), and maintained in microglial maintenance media for 9 days. 24-h before the assay, medium was changed to serum-free neuro-glial differentiation media^89^ to remove GPR34 ligand. Cells were dosed with either vehicle control or 25nM GPR34 agonist M1 for 30 min. Cells were washed with ice-cold TBS and lysed on ice with RIPA buffer supplemented with 1X HALT protease and phosphatase inhibitor (Thermofisher 1861281) and protein concentrations were measured using BCA assay. Equal protein amounts were prepared with 1X NuPAGE LDS sample buffer (Invitrogen, NP0007) and 1X NuPAGE reducing agent (Invitrogen, NP0009), then boiled for 5 min at 95°C and subjected to SDS PAGE in NuPAGE 4-12% Bis-Tris Mini Protein Gels (Invitrogen, NP0321). Protein was transferred to PVDF membrane (Millipore, IPFL85R) overnight by wet transfer. Membranes were incubated for 1 hr in blocking buffer (LICOR, 927-60001) and then overnight at 4°C in primary antibodies (Vinculin: CST, 13901, ERK: CST, 4696, p-ERK: CST, 4370S) prepared 1:1000 in diluent (LICOR, 927-65001). The following day membranes were washed 3 x 5 min in TBS-T and incubated for 1 hr with secondary antibodies (LICOR 926-32213 and 926-32212) prepared 1:10,000 in LICOR diluent. Membranes were washed 3 x 5 min in TBS-T prior to exposure on LICOR Odyssey CLx Imager. Image acquisition and densitometry analysis were performed with ImageStudio Software (Version 5.5.4).

#### Seahorse cellular respiration assays

Line 1 iMGs (20,000 cells/well) were seeded on PDL-coated 96-well Agilent Seahorse XF Cell Culture microplate (103799-100) in microglia maintenance media. For the fatty acid oxidation experiments, three days prior to assay, media was removed and replaced with substrate-limited media (SLM) comprised of XF DMEM, 1% FBS, 0.5 mM glucose, 1 mM glutamine, and 0.5 mM carnitine. Glutamine was subsequently removed from media for another 24 hours. On the day of the assay, cells were washed twice with assay media comprised of XF DMEM, 2 mM glucose, and 0.5 mM carnitine for the fatty acid oxidation experiments. Cells were imaged using brightfield microscopy to obtain cell counts utilized for normalization. Cells were then incubated for 1 hour in a non-CO2 incubator before beginning the Seahorse experiment. Ports on the sensor plate were filled according to the XF Long Chain Fatty Acid Oxidation Stress Kit (103693-100, Agilent) and cells were subjected to sequential injections of oligomycin (final concentration 1.5 uM), FCCP (2 uM for fatty acid oxidation, 1.5 uM for glucose oxidation), and rotenone/antimycin A (0.5 uM each). In experiments using inhibitors, etomoxir were added in port A at 4 uM final concentration, respectively. Data was analyzed using the Agilent Seahorse Analytics online software to generate kinetic curves and calculate maximal respiration and spare capacity.

### Mouse brain imaging

#### Immunohistochemistry

Fresh, PBS-perfused brain hemispheres were immersion fixed for 24 h in 4% PFA (Electron Microscopy Sciences, Hatfield, PA) in PBS before gelatin embedding and sectioning (40 µm) at Neuroscience Associates using MultiBrain technology (Neuroscience Associates, Knoxville, TN). NSA-embedded sections were photobleached for 48h in PBS using LED transilluminators at 4°C. After photobleaching, sections were permeabilized and blocked for 4h at RT in 0.03% TX100 (Millipore Sigma catalog no. T8787)/0.125% CD5 (heptakis(2,6-di-*O*-methyl)-β-cyclodextrin; Millipore Sigma catalog no. 39915-1G)/4% Normal Donkey Serum (NDS; Jackson Immunoresearch catalog no. 017-000-121)/4% Fatty Acid-Free BSA (FAF-BSA; Millipore Sigma A7030)/1x Fish Gelatin (bioWorld catalog no. 20720008-1) in PBS. Sections were washed in PBS and stained for 48h with designated primary antibodies in 4% normal donkey serum/4% FAF-BSA in PBS at 4C. This was followed by 3×5 min washes in PBS, and overnight incubation in secondary antibodies at 4C. Sections were incubated with DAPI in PBS for 20 min (0.1 µg/ml), followed by two washes in PBS, and mounted on 2” x 3” glass slides using ProLong Glass (Thermo Fisher, P36982). Immunofluorescence staining was performed with the following primary antibodies: guinea pig anti-Iba1 (Synaptic Systems, HS234308, 1:500), rat anti-CD68 (Bio-Rad, MCA1957, 1:500), biotinylated mouse anti-Amyloid β (IBL America, 10326, 1:500), rabbit anti-Amyloid β (IBL America, 18584, 1:500) and the following secondary antibodies: donkey anti-rat Alexa Flour 488 (Invitrogen, A21208, 1:800), donkey anti-rabbit Alexa Fluor 568 (Invitrogen, A10042, 1:800), donkey anti-guinea pig Alexa Fluor 647 (Jackson Immunoresearch, 706-605-148, 1: 800), Streptavidin conjugated Alexa Fluor 750 (Invitrogen, S21384, 1:800), and donkey anti-rabbit CF750 (Biotium, 20298, 1:800).

#### Imaging and analysis of plaque compaction

Images of mouse brain sections fluorescently stained for Amyloid β were captured using a Zeiss Axioscan.Z1 slide scanner with a 20x/0.8 NA objective. A custom macro in Zeiss ZEN software was used to quantify plaque area by using median smoothing then local background subtraction to create a binary mask for each plaque followed by splitting each 4-connected mask component into an individual plaque object. Objects smaller than 1 μm^2^ were excluded as staining artifacts/debris. For each section, the distribution of plaque areas was calculated using gaussian kernel density estimation using the “gaussian_kde” function in scipy (v1.15)^90^, sampled on a log spaced domain between 1 and 1000 μm^2^. Differences between plaque distributions between *App*^SAA^-KI:*Gpr34* WT and KO animals was assessed using a Kolmogorov-Smirnov test implemented with the “ks_2samp” function in scipy. The *Gpr34* KO plaque distribution was normalized to the APP distribution by subtracting the mean distribution of all *App*^SAA^:GPR34 WT animals from each KO animal, then calculating the new mean and standard error of the difference.

#### Imaging and analysis of microglia morphology

Mouse brain sections fluorescently stained for Iba1 and CD68 were imaged on a Leica SP8 laser scanning confocal using a 25x/0.95 NA objective in Lightning superresolution mode. A custom segmentation and morphology analysis pipeline was used to segment the Iba1+ microglia and analyze their morphology using a multi-metric approach as described previously^22^. Briefly, the Iba1 channel was down sampled to an isotropic 0.5 μm voxel size, then background subtracted by subtracting a gaussian smoothed version of the channel with a kernel size of 10 μm. Individual microglia were split into 8-connected objects then any object larger than ∼10,000 μm^3^ was split into approximately equal sized chunks using k-means clustering where k = ceiling(volume / 10,000 μm^3^). Microglia fragments smaller than 500 μm^3^ were excluded from downstream analysis. For each microglia, the object was morphologically eroded by 1 μm^3^, removing all thin processes, and the resulting mask was defined as the microglial “core”. The inverse of this mask was then defined as the “surface”. The morphological skeleton of each object was calculated using the “skeletonize” function in scikit-image (v0.25)^91^ and then continuous branches extracted by removing all skeleton voxels with >2 connectivity to the rest of the skeleton. Mean expression of Iba1 and CD68 was calculated inside the whole microglia mask as well as in the core and surface compartments for each microglia. To determine signature features of each genotype, morphological and intensity feature differences per condition were ranked by both effect size (Cohen’s d) and by FDR corrected t-tests using the functions in statsmodels (v0.14)^92^ with one-vs-rest comparisons. Four consistently top-ranking features were selected for the final time course analysis.

#### Quantification and statistical analyses

Unless otherwise indicated, statistical analyses were performed using GraphPad Prism Software (Version 10.2.2, 341) or R Studio. Statistical details including specific tests used, exact value of n, e.g. number of animals, cells, or independent trials, and the definition of center, dispersion, and precision measures e.g. mean, median, SD, SEM, are indicated in the corresponding figure legend. All in vitro data are from at least three independent experiments. All statistical analyses performed were two-tailed. If data met statistical assumptions, unpaired *t*-test, paired *t*-test, Sidak’s multiple comparisons test, or Tukey’s multiple comparisons test was used depending on data groups. Statistical *p* values are shown on graphs denoted with asterisks, \**p*<0.05, \*\**p*<0.01, \*\*\**p*<0.001, ns: not significant.

## KEY RESOURCES TABLE

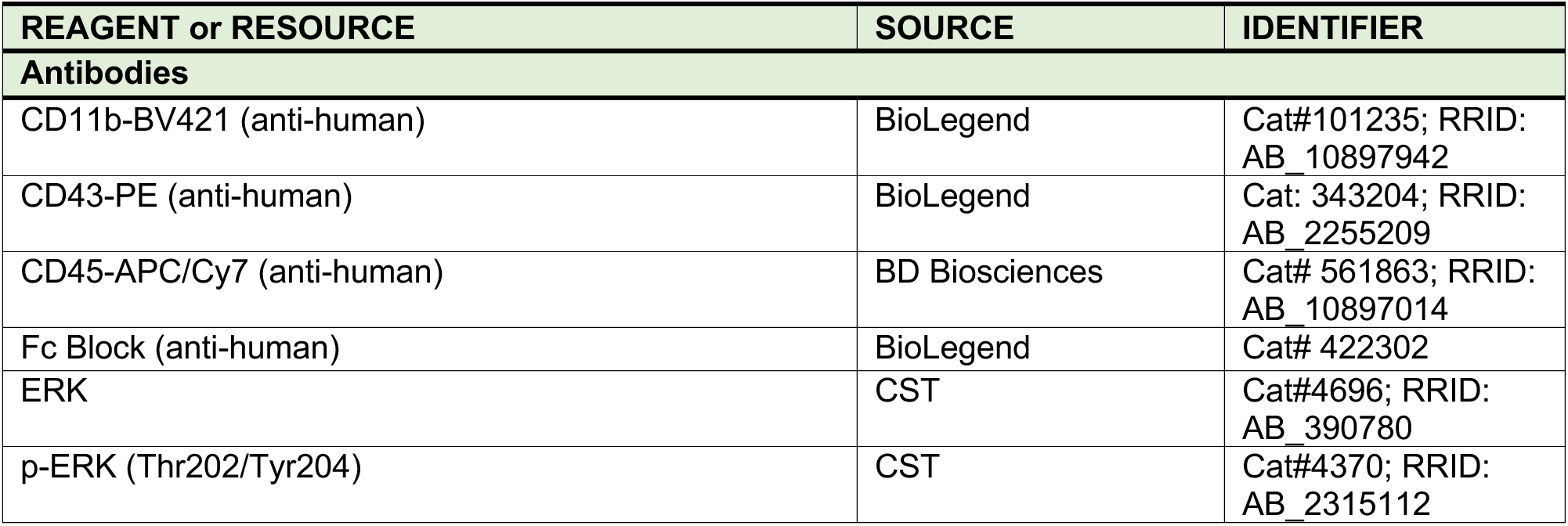
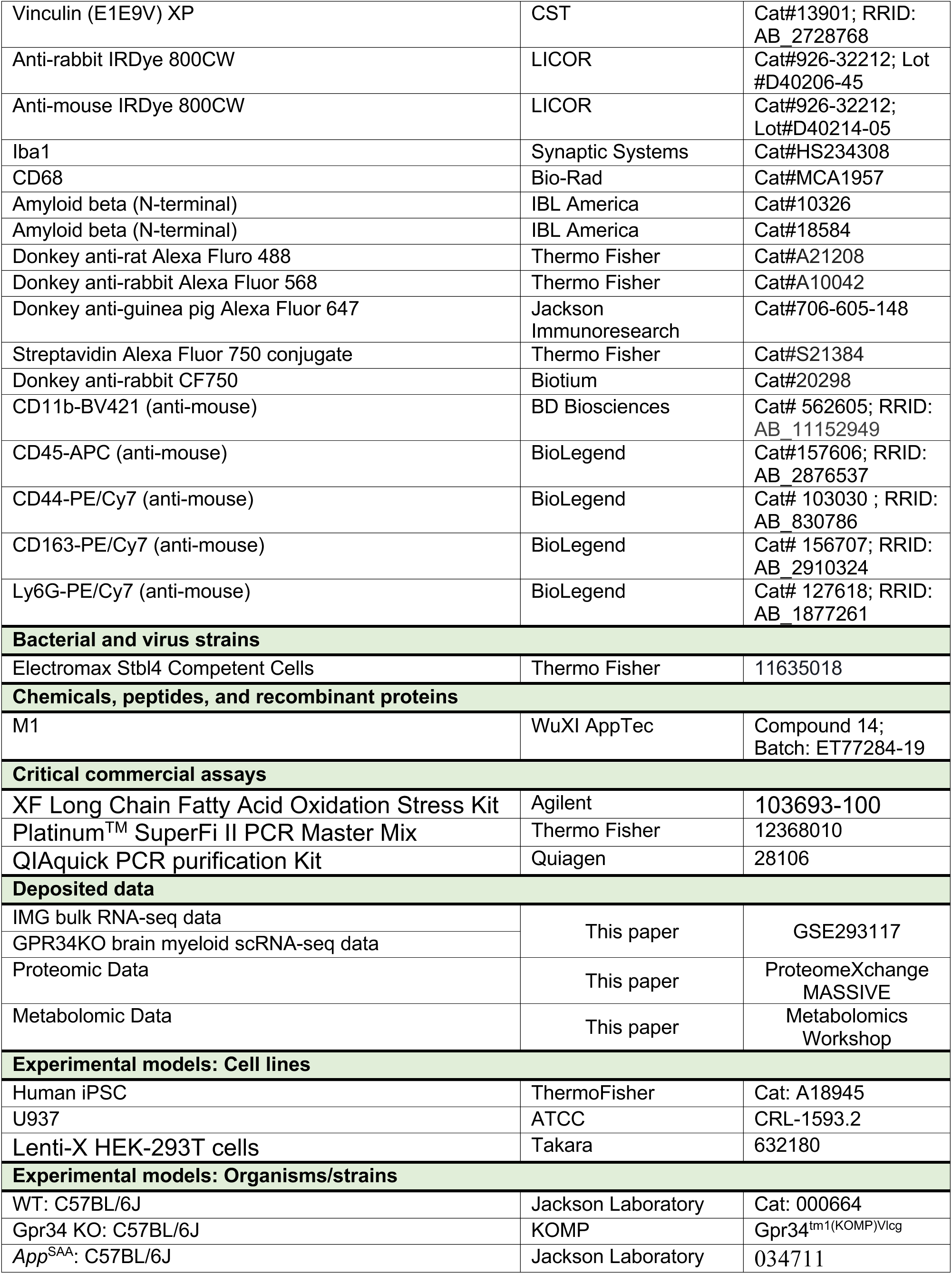
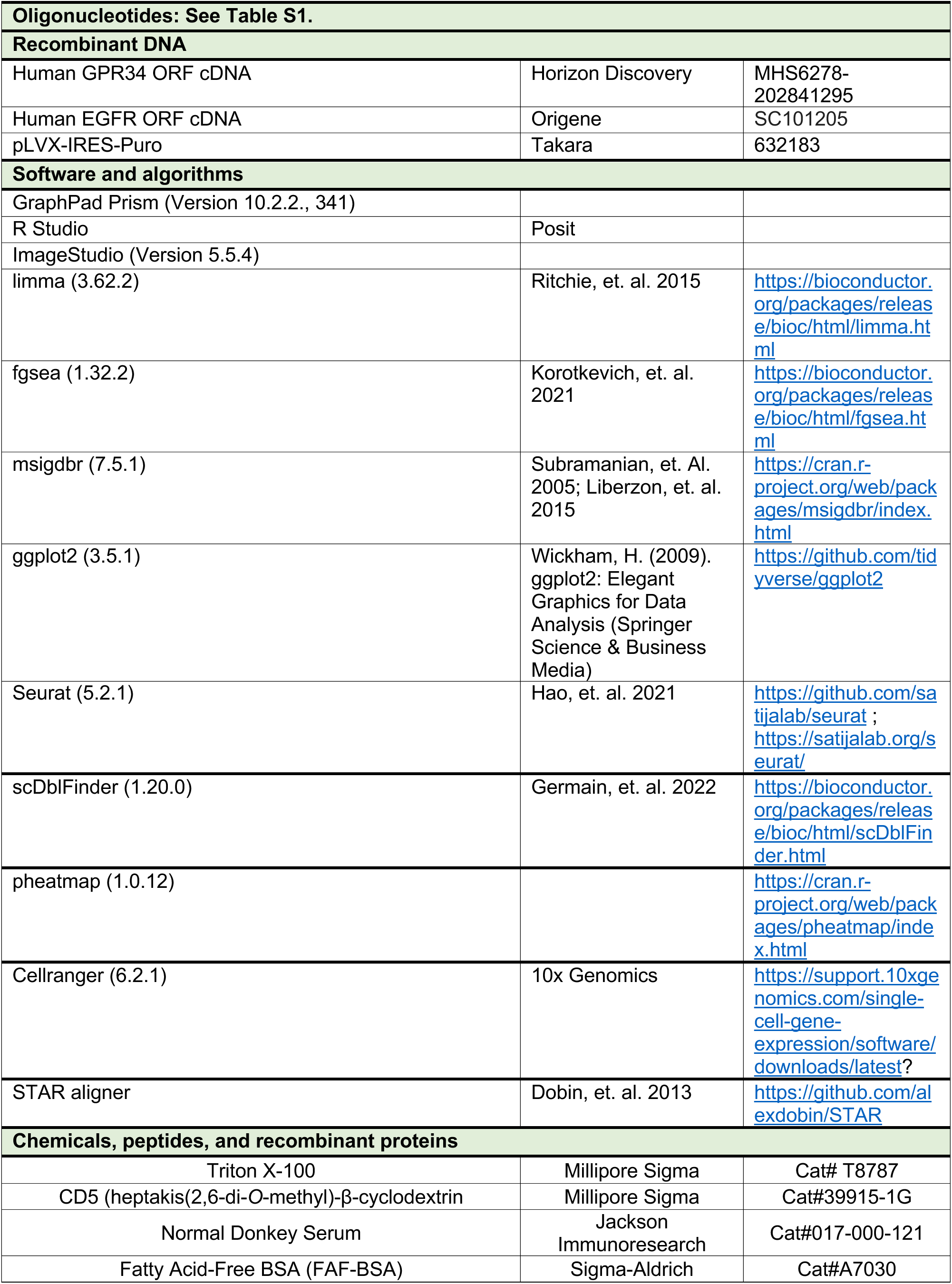
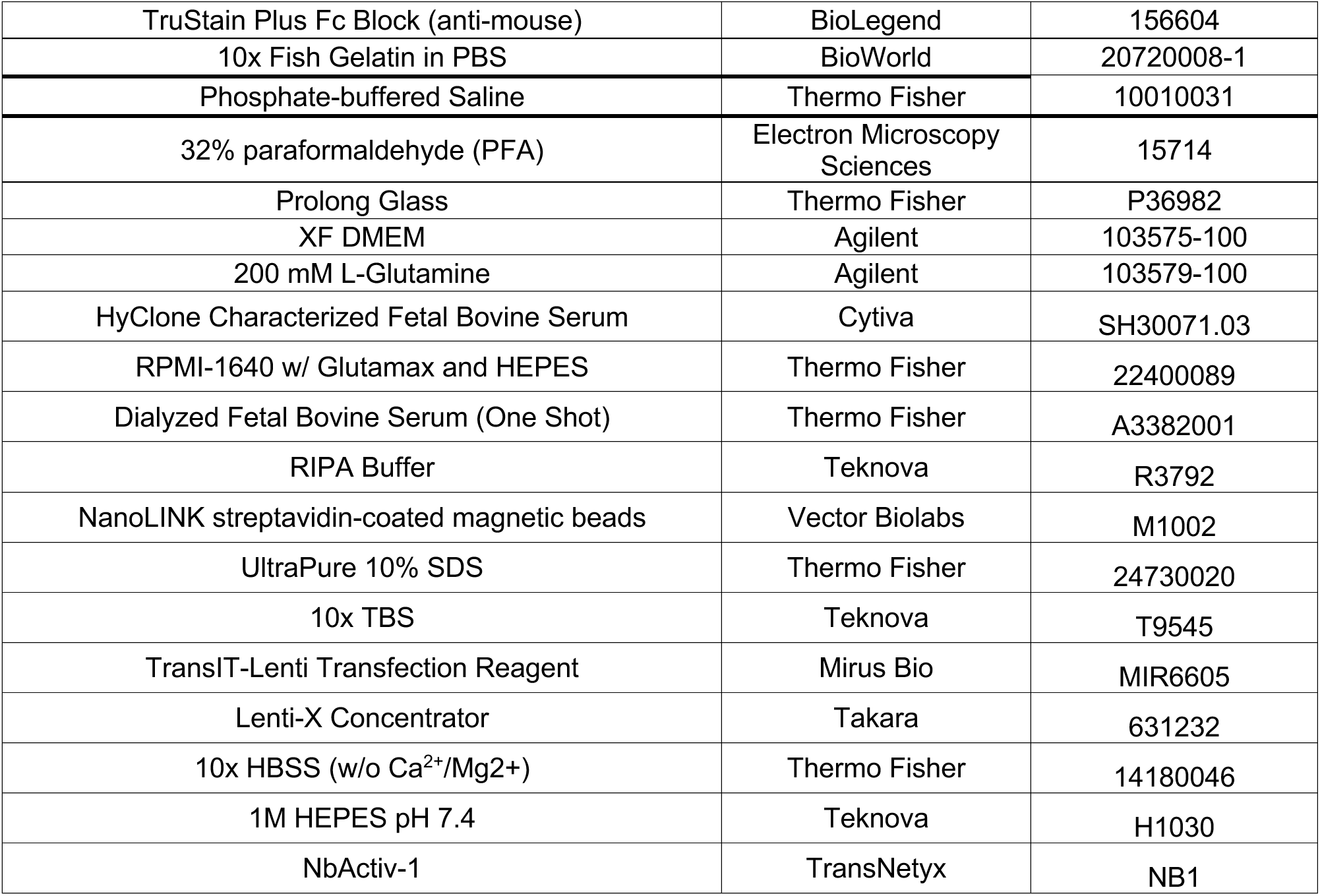

