## Supplementary material for "GPR34 regulates microglia state and loss-of-function rescues TREM2 metabolic dysfunction": Figure S1 to S4

**A**

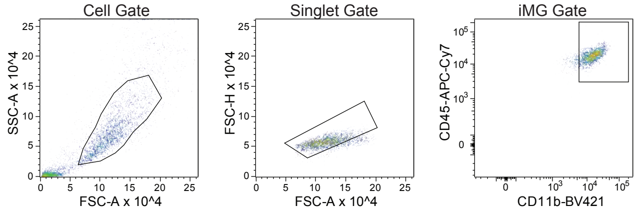


**B**

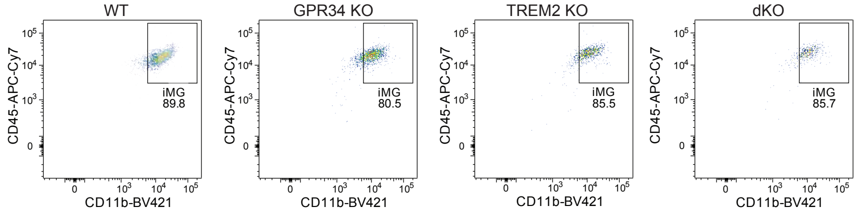


**C**

#### gRNA sequences

**GPR34** CUAGGUGUGAUUCUGUGCA
**TREM2** TATGACTCCATGAAGCACTG

**D**

#### gRNA sequences

**AAVS1-1** CCAUCGUAAGCAAACCUUAGGUUUUAGAGCUAUGCU
**AAVS1-2** GGGACCACCUUAUAUUCCCAGUUUUAGAGCUAUGCU
**AAVS1-3** GACGCAAGGGAGACAUCCGUGUUUUAGAGCUAUGCU

**GPR34-1** UCGGUCUAUACAGCAACGGAGUUUUAGAGCUAGAAAUAGCAAGUUAAAAUAAGGCUAGUCCGUUAUCAACUUGAAAAAGUGGCACCGAGUCGGUGCUUUU
**GPR34-2** AUUUUCAUCGUGGGACUGGUGUUUUAGAGCUAGAAAUAGCAAGUUAAAAUAAGGCUAGUCCGUUAUCAACUUGAAAAAGUGGCACCGAGUCGGUGCUUUU
**GPR34-3** GGUUAUAAAGCGCAUUCUGUGUUUUAGAGCUAGAAAUAGCAAGUUAAAAUAAGGCUAGUCCGUUAUCAACUUGAAAAGUGGCACCGAGUCGGUGCUUUU

**TREM2-1** UGCUGACCACACGCUGGCAU
**TREM2-2** CUGGAACACUGUGGUGUUGU
**TREM2-3** CCUAUGACUCCAUGAAGCAC

**E**

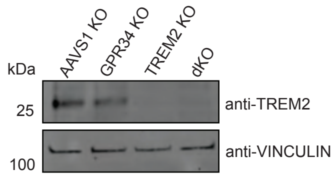


**F**

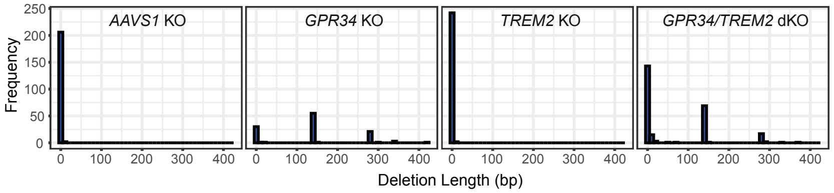


**G**

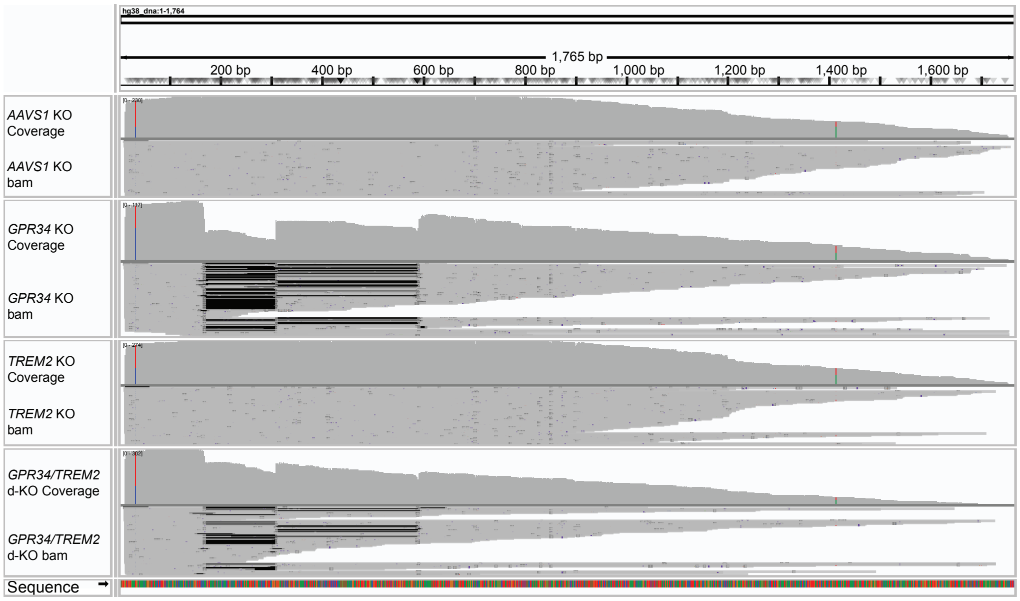


### Figure S1

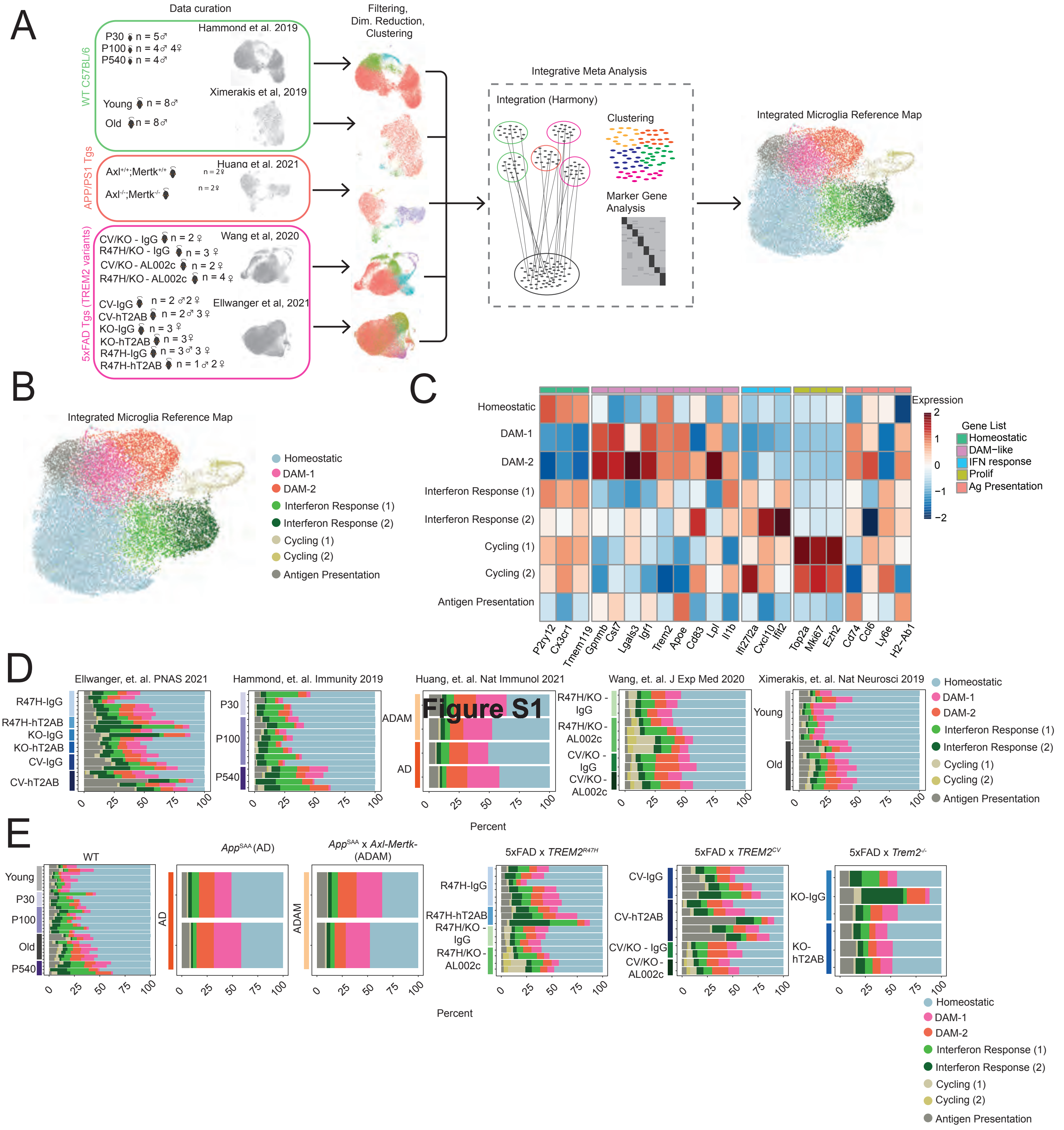
A
Data curation
Filtering, Dim. Reduction, Clustering
WT C57BL/6
Hammond et al. 2019
P30 n = 5♂
P100 n = 4♂ 4♀
P540 n = 4♂
Ximerakis et al, 2019
Young n = 8♂
Old n = 8♂
APP/PS1 Tgs
Huang et al. 2021
Axl+/+;Mertk+/+ n = 2♀
Axl-/-;Mertk-/- n = 2♀
5xFAD Tgs (TREM2 variants)
Wang et al, 2020
CV/KO - IgG n = 2♀
R47H/KO - IgG n = 3♀
CV/KO - AL002c n = 2♀
R47H/KO - AL002c n = 4♀
Ellwanger et al, 2021
CV-IgG n = 2♂ 2♀
CV-hT2AB n = 2♂ 3♀
KO-IgG n = 3♀
KO-hT2AB n = 3♀
R47H-IgG n = 3♂ 3♀
R47H-hT2AB n = 1♂ 2♀
Integrative Meta Analysis
Integration (Harmony)
Clustering
Marker Gene Analysis
Integrated Microglia Reference Map
B
Integrated Microglia Reference Map
Homeostatic
DAM-1
DAM-2
Interferon Response (1)
Interferon Response (2)
Cycling (1)
Cycling (2)
Antigen Presentation
C
Expression
Gene List
Homeostatic
DAM−like
IFN response
Prolif
Ag Presentation
P2ry12 Cx3cr1 Tmem119 Gpnmb Cst7 Lgals3 Igf1 Trem2 Apoe Cd83 Lpl Il1b Ifi27l2a Cxcl10 Ifit2 Top2a Mki67 Ezh2 Cd74 Ccl6 Ly6e H2-Ab1
D
Ellwanger, et. al. PNAS 2021
Hammond, et. al. Immunity 2019
Huang, et. al. Nat Immunol 2021
Wang, et. al. J Exp Med 2020
Ximerakis, et. al. Nat Neurosci 2019
R47H-IgG
R47H-hT2AB
KO-IgG
KO-hT2AB
CV-IgG
CV-hT2AB
P30
P100
P540
ADAM
AD
Figure S1
R47H/KO - IgG
R47H/KO - AL002c
CV/KO - IgG
CV/KO - AL002c
Young
Old
Percent
E
WT
App^SAA (AD)
App^SAA x Axl-Mertk- (ADAM)
5xFAD x TREM2^R47H
5xFAD x TREM2^CV
5xFAD x Trem2-/-
Percent

**Figure S2**

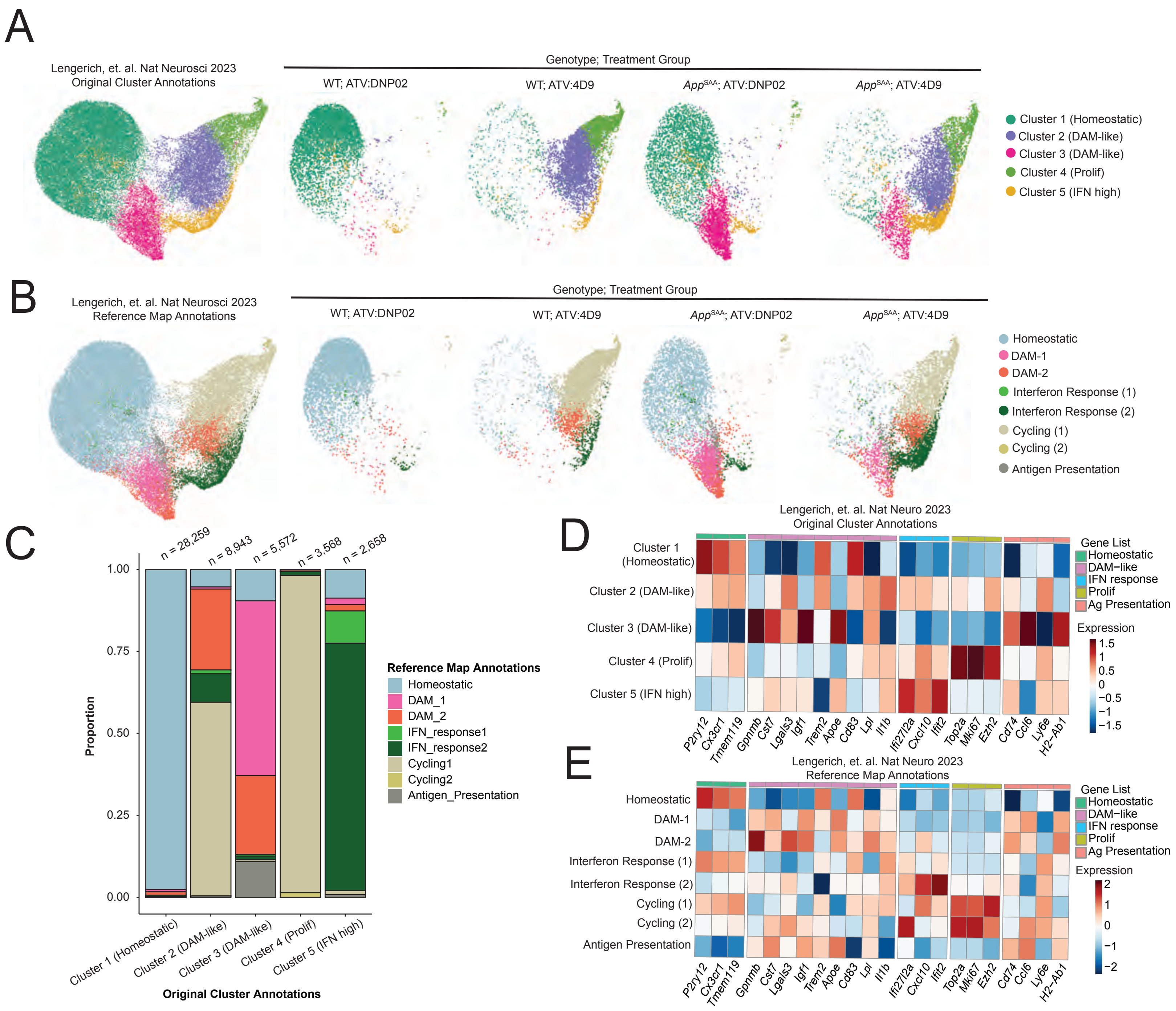
A
Lengerich, et. al. Nat Neurosci 2023
Original Cluster Annotations
Genotype; Treatment Group
WT; ATV:DNP02
WT; ATV:4D9
App^SAA; ATV:DNP02
App^SAA; ATV:4D9
Cluster 1 (Homeostatic)
Cluster 2 (DAM-like)
Cluster 3 (DAM-like)
Cluster 4 (Prolif)
Cluster 5 (IFN high)
B
Lengerich, et. al. Nat Neurosci 2023
Reference Map Annotations
Genotype; Treatment Group
WT; ATV:DNP02
WT; ATV:4D9
App^SAA; ATV:DNP02
App^SAA; ATV:4D9
Homeostatic
DAM-1
DAM-2
Interferon Response (1)
Interferon Response (2)
Cycling (1)
Cycling (2)
Antigen Presentation
C
n = 28,259
n = 8,943
n = 5,572
n = 3,568
n = 2,658
1.00
0.75
0.50
0.25
0.00
Proportion
Cluster 1 (Homeostatic)
Cluster 2 (DAM-like)
Cluster 3 (DAM-like)
Cluster 4 (Prolif)
Cluster 5 (IFN high)
Original Cluster Annotations
Reference Map Annotations
Homeostatic
DAM_1
DAM_2
IFN_response1
IFN_response2
Cycling1
Cycling2
Antigen_Presentation
D
Lengerich, et. al. Nat Neuro 2023
Original Cluster Annotations
Cluster 1 (Homeostatic)
Cluster 2 (DAM-like)
Cluster 3 (DAM-like)
Cluster 4 (Prolif)
Cluster 5 (IFN high)
P2ry12 Cx3cr1 Tmem119 Gpnmb Cst7 Lgals3 Igf1 Trem2 Apoe Cd83 Lpl Il1b Ifi27l2a Cxcl10 Ifit2 Top2a Mki67 Ezh2 Cd74 Ccl6 Ly6e H2-Ab1
Gene List
Homeostatic
DAM−like
IFN response
Prolif
Ag Presentation
Expression
1.5
1
0.5
0
−0.5
−1
−1.5
E
Lengerich, et. al. Nat Neuro 2023
Reference Map Annotations
Homeostatic
DAM-1
DAM-2
Interferon Response (1)
Interferon Response (2)
Cycling (1)
Cycling (2)
Antigen Presentation
P2ry12 Cx3cr1 Tmem119 Gpnmb Cst7 Lgals3 Igf1 Trem2 Apoe Cd83 Lpl Il1b Ifi27l2a Cxcl10 Ifit2 Top2a Mki67 Ezh2 Cd74 Ccl6 Ly6e H2-Ab1
Gene List
Homeostatic
DAM−like
IFN response
Prolif
Ag Presentation
Expression
2
1
0
−1
−2

**Figure S3**

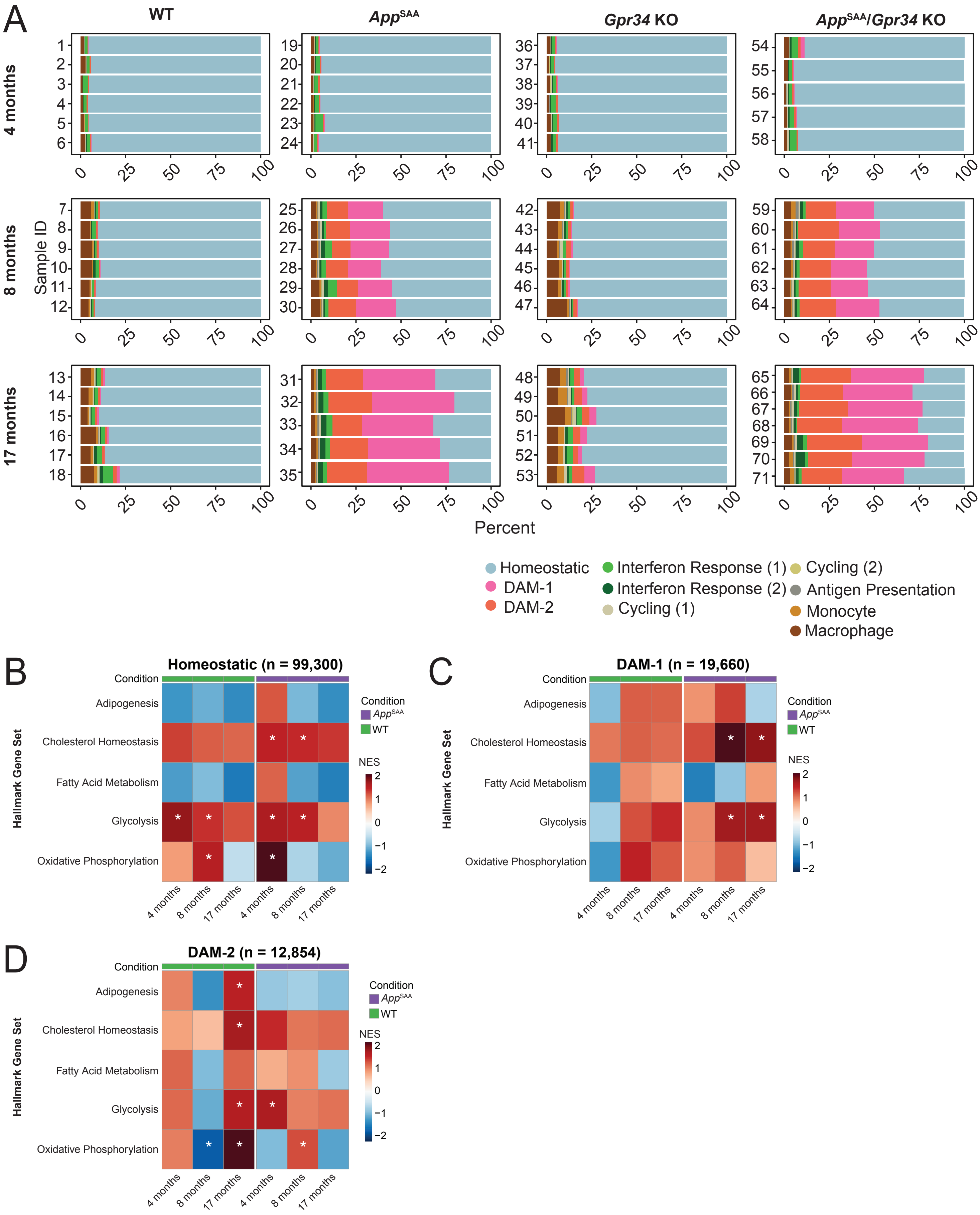
A
WT
AppSAA
Gpr34 KO
AppSAA/Gpr34 KO
4 months
8 months
17 months
Sample ID
Percent
0
25
50
75
100
Homeostatic
DAM-1
DAM-2
Interferon Response (1)
Interferon Response (2)
Cycling (1)
Cycling (2)
Antigen Presentation
Monocyte
Macrophage
B
Homeostatic (n = 99,300)
C
DAM-1 (n = 19,660)
D
DAM-2 (n = 12,854)
Condition
Hallmark Gene Set
Adipogenesis
Cholesterol Homeostasis
Fatty Acid Metabolism
Glycolysis
Oxidative Phosphorylation
AppSAA
WT
NES
2
1
0
−1
−2
4 months
8 months
17 months

Figure S4
