## Supplementary figures and images for "GPR34 regulates microglia state and loss-of-function rescues TREM2 metabolic dysfunction"

### Figure S5

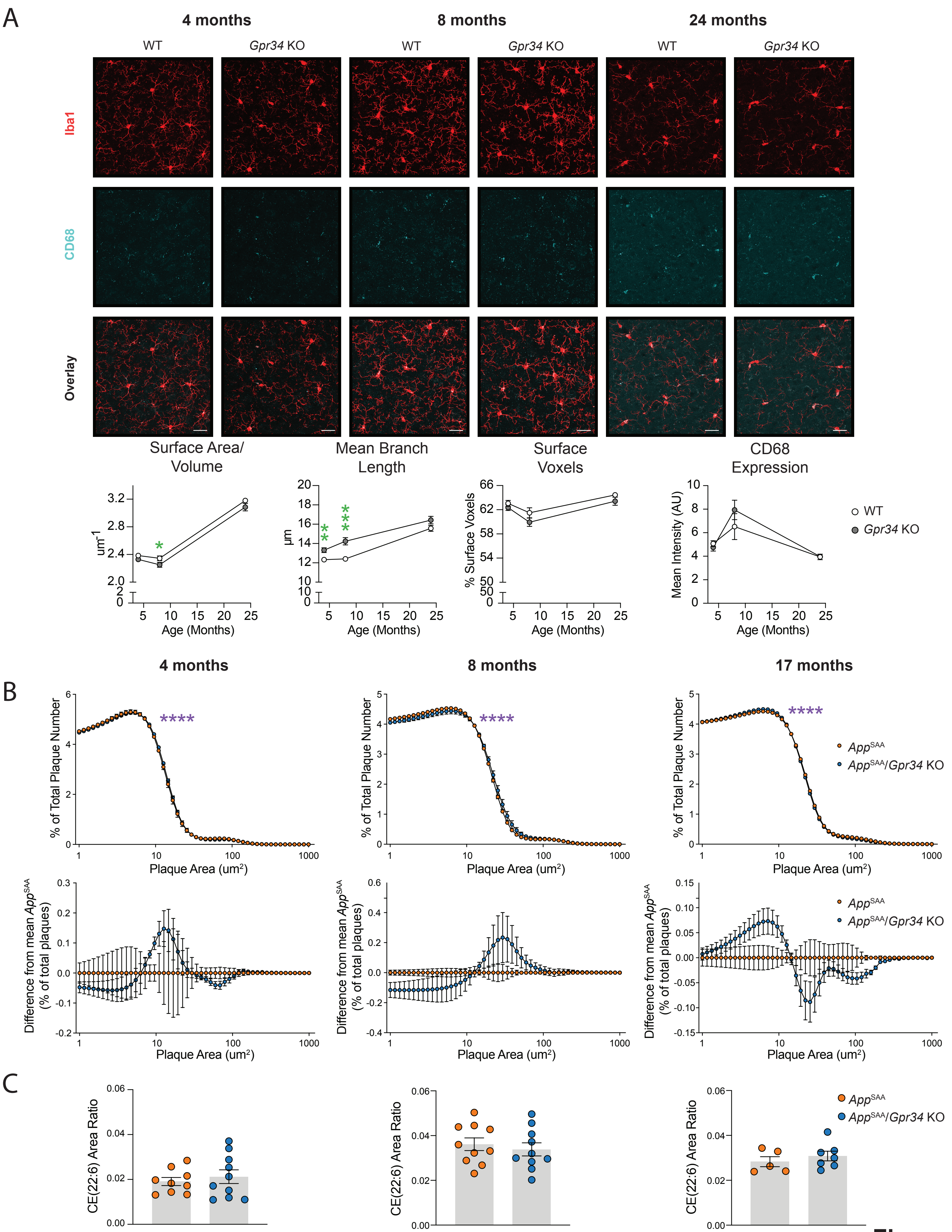
